# Integrated multi-omics highlights alterations of gut microbiome functions in prodromal and idiopathic Parkinson’s disease

**DOI:** 10.1101/2024.12.13.628341

**Authors:** Remy Villette, Júlia Ortís Sunyer, Polina V. Novikova, Velma T. E. Aho, Viacheslav A. Petrov, Oskar Hickl, Susheel Bhanu Busi, Charlotte De Rudder, Benoit Kunath, Anna Heintz-Buschart, Jean-Pierre Trezzi, Rashi Halder, Christian Jäger, Laura A. Lebrun, Annegrät Daujeumont, Sebastian Schade, Annette Janzen, Nico Jehmlich, Martin von Bergen, Cédric C. Laczny, Patrick May, Claudia Trenkwalder, Wolfgang Oertel, Brit Mollenhauer, Paul Wilmes

## Abstract

Parkinson’s disease (PD) is associated with gut microbiome shifts, but the functional consequences remain unclear. Here, we use an integrated multi-omics approach to compare the gut microbiomes of individuals with PD and prodromal PD as well as healthy individuals. After analyzing each omics, meta-metabolomic was selected to inform the analysis as it represents the most discriminatory and robust ome. We identified 11 metabolites that were differentially abundant between the groups, amongst which β-glutamate was increased in PD and prodromal PD, and correlated with the transcriptional activities of *Methanobrevibacter smithii* and *Clostridium* spp. We identified decreases in transcripts, but not in gene abundances, related to glutamate metabolism, bile acids, chemotaxis and flagellar assembly in PD, particularly in keystone genera such as *Roseburia, Agathobacter* and *Blautia*. Our findings, integrated into the Expobiome map, reveal multifactorial microbiome alterations which converge with PD pathways. Our study highlights the importance of investigating the gut microbiome’s functional dimensions to better resolve microbiome-host interactions in health and disease.

## Introduction

Parkinson’s disease (PD), a neurodegenerative disease impacting movement due to dopaminergic neuron loss, is the second most prevalent neurodegenerative disease worldwide^1^. Individuals with PD are often characterized by an increase in gut permeability, inflammation and constipation which, together, suggest a link between the gut microbiome and PD etiology^2–4^. This potential link is supported by numerous studies reporting differences in the gut microbiome structure of individuals with PD compared to healthy individuals^5–11^. These findings have been further confirmed by recent meta-analyses^12,13^. Together, the studies highlight a decreased abundance for the genera *Roseburia*, *Blautia, Butyricoccus* and *Faecalibacterium* in PD while *Methanobrevibacter*, *Akkermansia*, *Lactobacillus*, *Bifidobacterium* and *Hungatella*, are typically enriched^5–13^. Similar changes in idiopathic REM sleep behavior disorder (iRBD), a prodromal stage of PD^14^, have been reported^7,10^. Moreover, the taxa decreased in PD are known producers of short-chain fatty acids (SCFAs), which correspondingly have also been found to be decreased in concentration in fecal samples of PD individuals^6,15,16^.

In addition to SCFAs, several microbiome-derived metabolites such as bile acids (BAs), glycine and glutamate have been associated with PD, either in plasma, serum^17,18^ or stool^19–21^. BAs are produced by the host and metabolized by the gut microbiome into secondary bile acids with different cytotoxic capacities but also immunomodulatory capacities^22,23^. Glutamate is the major excitatory neurotransmitter and exerts toxic activity on neuronal cells^24^. Its levels in serum and cerebrospinal fluid have been reported as either increased^25,26^, not different^27^, or decreased^28^, but decreased in the gut in PD compared to healthy controls (HC)^29^.

Altogether, alterations of the gut microbiome are linked to PD, but less is known about iRBD or other prodromal stages of the disease. Moreover, most of the associations between PD and the gut microbiome are based on taxonomic and metabolomic analyses. The resulting data, although insightful, lacks functional and systemic information that could better capture the complex crosstalk between the gut microbiome and the host in the context of PD. To obtain such information, we performed an integrated multi-omics study on a cross-sectional cohort comprised of individuals with iRBD and PD alongside HC. Metagenomics (MG), metatranscriptomics (MT), metaproteomics (MP) and meta-metabolomics (MM) were used to characterize taxonomic (taxMG, taxMT, taxMP) and functional (funMG, funMT, funMP, MM) differences between HC, iRBD and PD gut microbiomes. We identified substantial differences in gut microbiome functions and metabolites between the groups, including an increase in β-glutamate levels in PD that were related to a dysregulation of glutamate-related gene expression. Alterations in glutamate-related genes were linked with chemotaxis and flagellar assembly pathways, for which we identified strong and distinct taxonomic differences in transcription between PD and HC. Collectively, our data highlight the importance of multi-omics approaches for the identification of microbiome-mediated effects on neurological, and more broadly, complex human diseases involving host-microbiome interactions.

## Results

### Study cohort

Our initial set of subjects consisted of 50 individuals with PD, diagnosed according to United Kingdom Parkinson’s Disease Society Brain Bank (UKPDSBB) clinical diagnostic criteria^30^, and 30 people with polysomnography-confirmed iRBD as well as 50 healthy control (HC) subjects. The data from 4 PD and 3 iRBD as well as 1 HC were subsequently excluded (see Methods), leading to a final data set of 46 PD, 27 iRBD and 49 HCs. The subjects in the three groups were of similar age but had slightly different gender distributions, with males overrepresented in the iRBD and PD groups (Table 1, Fisher test, p = 0.004), as is typical for these conditions^31,32^. Constipation, a prevalent non-motor symptom of PD^33^, was also more common in the iRBD and PD groups compared to HC (Fisher test, p < 0.001).

### Microbiome function is altered in PD

Alpha diversity comparisons revealed no statistically significant differences between the three groups when considering taxMG, taxMT, funMG, taxMP and MM (Fig. 2A, Mann-Whitney test, p > 0.05). However, funMT and funMP showed a statistically significant increase in alpha diversity in PD compared to HC and iRBD compared to PD, respectively (Figure 2A, Mann-Whitney test, p < 0.05). We then analyzed beta diversity for all omics layers. TaxMG, funMG and taxMP revealed no statistically significant differences between the three groups (Extended fig. 1 A, B and D, PERMANOVA, p= 0.2, 0.8 and 0.35, respectively), while taxMT, funMT, funMP and MM showed a statistically significant separation of the groups, especially for funMT (Figure 2B and C, Extended fig. 1 C, PERMANOVA p = 0.001, 0.001, 0.005 and p = 0.008, respectively). Permutation analysis revealed that funMT, funMP and MM resulted in the best separation of the three groups while taxMG, funMG and taxMP exhibited the lowest separation capacity (Extended fig. 1D, PERMANOVA R² of 0.043, 0.041, 0.038, 0.018, 0.014 and 0.017, respectively). Pairwise comparisons using PERMANOVA demonstrated that most differences were found between either HC and PD or HC and iRBD, with only funMP and funMT showing statistically significant differences between PD and iRBD (Figure 2E). Based on these differences, we next assessed how the confounding factors sex, age and constipation may impact beta diversity. Sex and constipation were found to have a significant association with beta diversity for taxMT and funMT, while age was associated with taxMG (Figure 2F). Importantly, MM and funMP were not found to be associated with any confounders (Figure 2F).

**Figure 1.**
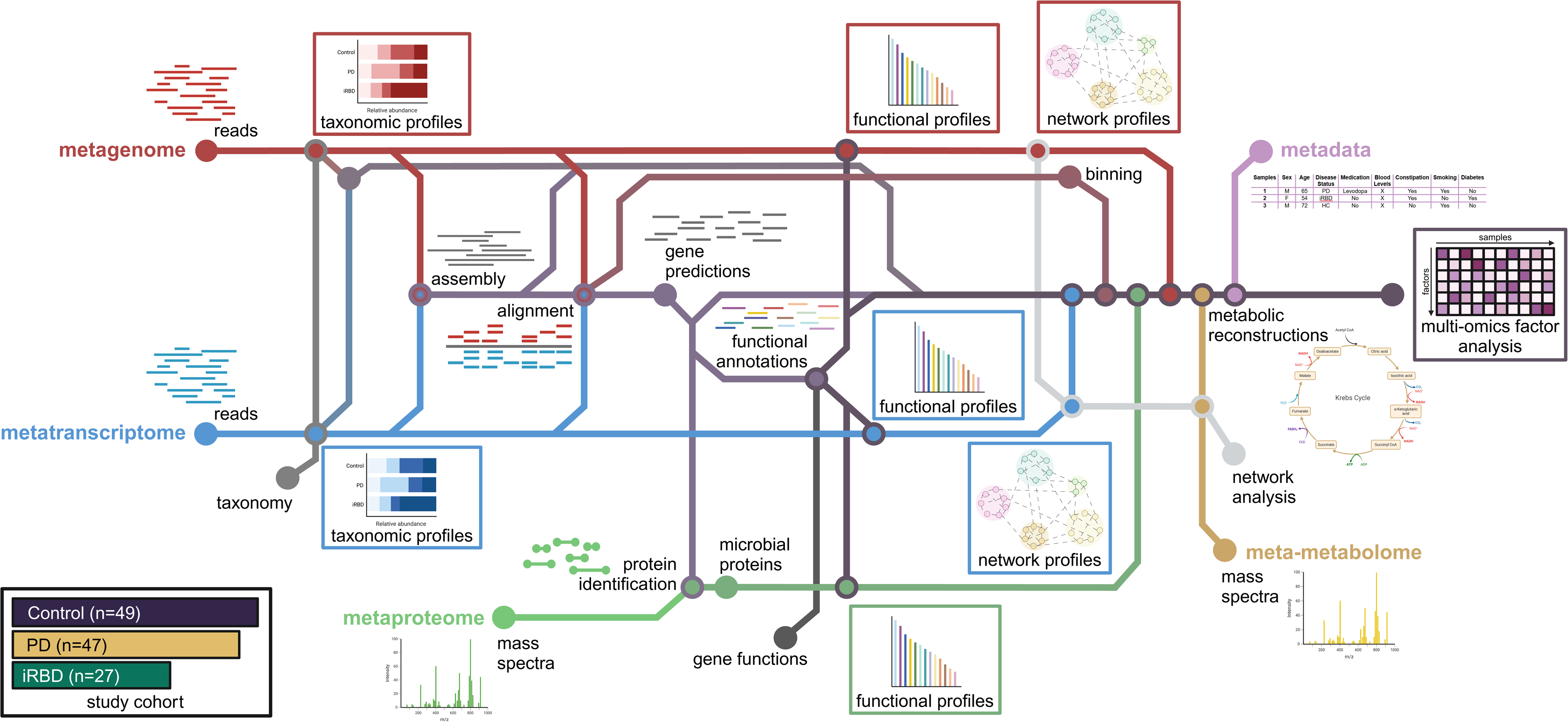
Schematic representation of the analytical workflow. Metagenomic (MG), metatranscriptomic (MT), metaproteomic (MP) and meta-metabolomic (MM) data were generated for each sample. Pre-processed MG and MT reads were sample-wise assembled using the iterative hybrid assembly pipeline of the Integrated Meta-omics Pipeline (IMP). After assembly, taxonomic annotation was performed at the read and contig levels, followed by gene prediction and functional annotation on the assembled contigs. Expressed proteins (MP) were identified using the predicted genes from the MG/MT hybrid assembly. For these three omics levels, we generated taxonomic and functional profiles that are referred to as taxMG, taxMT and taxMP for the taxonomic level, and funMG, funMT and funMP for the functional level, respectively. Community-based networks were reconstructed from gene annotations. Finally, the meta-metabolome (MM) was integrated with the other omics data at the network level. The integrated multi-omics analysis was performed with the available clinical metadata.

**Figure 2.**
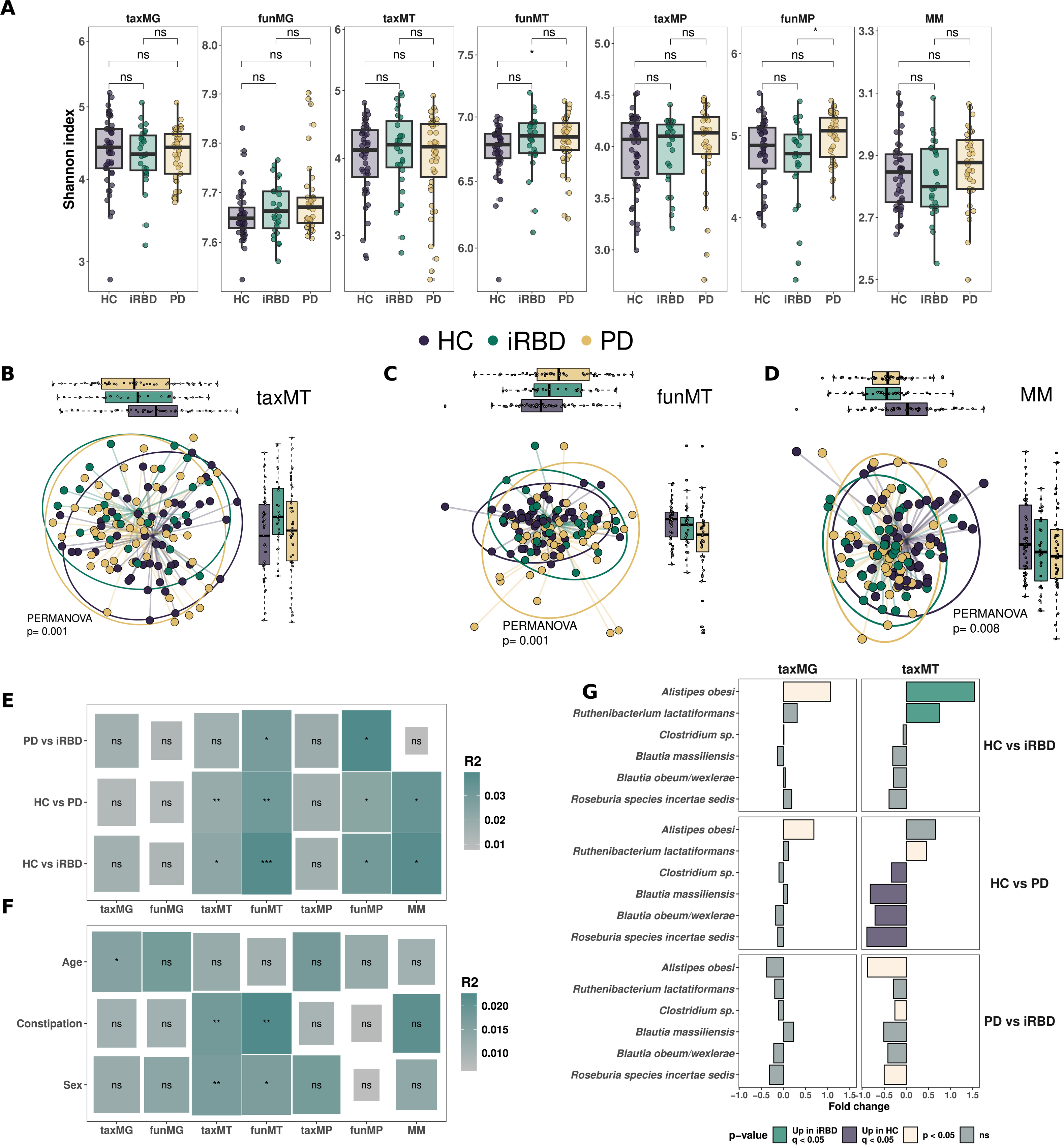
Microbiome structure is altered in PD and iRBD vs HC. **A.** Shannon index for different omics between Healthy Controls (HC), idiopathic REM sleep behaviour disorder (iRBD) and Parkinson’s Disease (PD). P-values are based on pairwise Mann-Whitney tests. Non-metric Multidimensional Scaling (NMDS) based on Bray-Curtis dissimilarity for taxonomic annotation of metatranscriptomic (taxMT) (**B**) and functional metatranscriptomic (funMT) data (**C**). **D**. Principal Component Analysis (PCA) for meta-metabolomic (MM) data based on untargeted, targeted SCFA and targeted bile acids abundance. All three quantifications have been sum-normalised before any merging. PCA was then computed on the merged matrix. All tests are based on PERMANOVA with 1000 permutations. **E**. Pairwise PERMANOVA between groups for each omics. **F**. PERMANOVA analysis for “Age”, “Sex” and “Constipation” for each omics. Size of rectangle is based on – log10(p-value) and colour on R² value. **G**. Differential abundance analysis using SIAMCAT for taxMG and taxMT. Values are pseudo fold changes for pairwise comparison between groups and size is based on –log10(p-value). Shape is based on significance before and after FDR correction.

We next looked at differential abundances on taxonomic layers to highlight the compositional differences between the groups. No significant differences were found in taxMG at the genus and species levels between any of the three groups (Figure 2G and Extended fig. 2C, q > 0.05, SIAMCAT). For taxMT, SIAMCAT highlighted an increase in *Alistipes obesi* and *Ruthenibacterium lactatiformans* in iRBD vs HC, *Roseburia incertae sedis*, *Blautia massiliensis, B. obeum* and *Clostridium* sp. were decreased in PD vs HC and no significant differences were found between PD and iRBD (Figure 2G, q < 0.05, q < 0.05 and q > 0.05, respectively). We identified the genus *Eubacterium* as being decreased in PD compared to HC in taxMT (Extended fig. 2C, q < 0.05, SIAMCAT). The ALDEx2 algorithm highlighted only *Roseburia incertae sedis* to be depleted in PD after FDR correction (Extended fig. 2B).

Subsequently, we investigated overall differences in gene abundances and expressions of the microbial genes linked to the observed differences in metabolites using the KEGG database. We observed no statistically significant differences in gene abundances between HC vs PD or HC vs iRBD (funMG, Extended fig. 4A and B, q > 0.05). Gene expression highlighted one transcript upregulated in HC vs iRBD, but 145 transcripts upregulated in HC vs PD (funMT, Extended fig. 4C and D, q < 0.05 and q > 0.05, respectively). No genes or transcripts were significantly higher in iRBD or PD compared to HC (Extended fig. 4A-D, q > 0.05). Finally, we found no statistically significant differences in gene or transcript abundances between PD and iRBD (data not shown).

### Altered metabolome of PD patients is linked to microbial abundance and activity

Considering that MM is associated with PD and iRBD but not with confounders, we chose MM as a robust guide for further statistical comparisons. In addition, MM can be considered as one of the final outputs of microbial activity and an important driver of microbial effects on the host. For this purpose, we removed unidentified compounds from the statistical testing, because we cannot associate microbial genes with them. Our analyses revealed 11 statistically significantly different compounds between the groups, including alanine, β-glutamate, serine, and glycerol (Figure 3A, q < 0.05). We found a significant increase in isovalerate, isobutyrate and valerate in PD patients (Figure 3A, q < 0.05) but no differences for butyrate, acetate, formate, propionate or total SCFAs (data not shown). Primary bile acids glycocholic and chenodeoxycholic acids were decreased in PD and in iRBD and PD, respectively (Figure 3A, q < 0.05). Glutamate was not differentially abundant between the three groups (data not shown). To unravel the effect of confounding factors, we measured the variance explained by each factor on the compounds’ abundances. Diagnosis explained more variance than constipation or sex (Fig. 3B, diagnosis: mean= 4.18%; median = 3.33%, sex: mean = 2.08%, median = 1.3%; constipation: mean = 3.7%, median = 2.68%). We found only malic acid as being significantly associated with sex and no compound to be associated with constipation (Extended fig. 3A and B, q < 0.05, q > 0.05). Based on the PERMANOVA results and variance analysis, the differences observed in MM were most strongly associated with the disease status and, importantly, not with the confounding factors.

**Figure 3.**
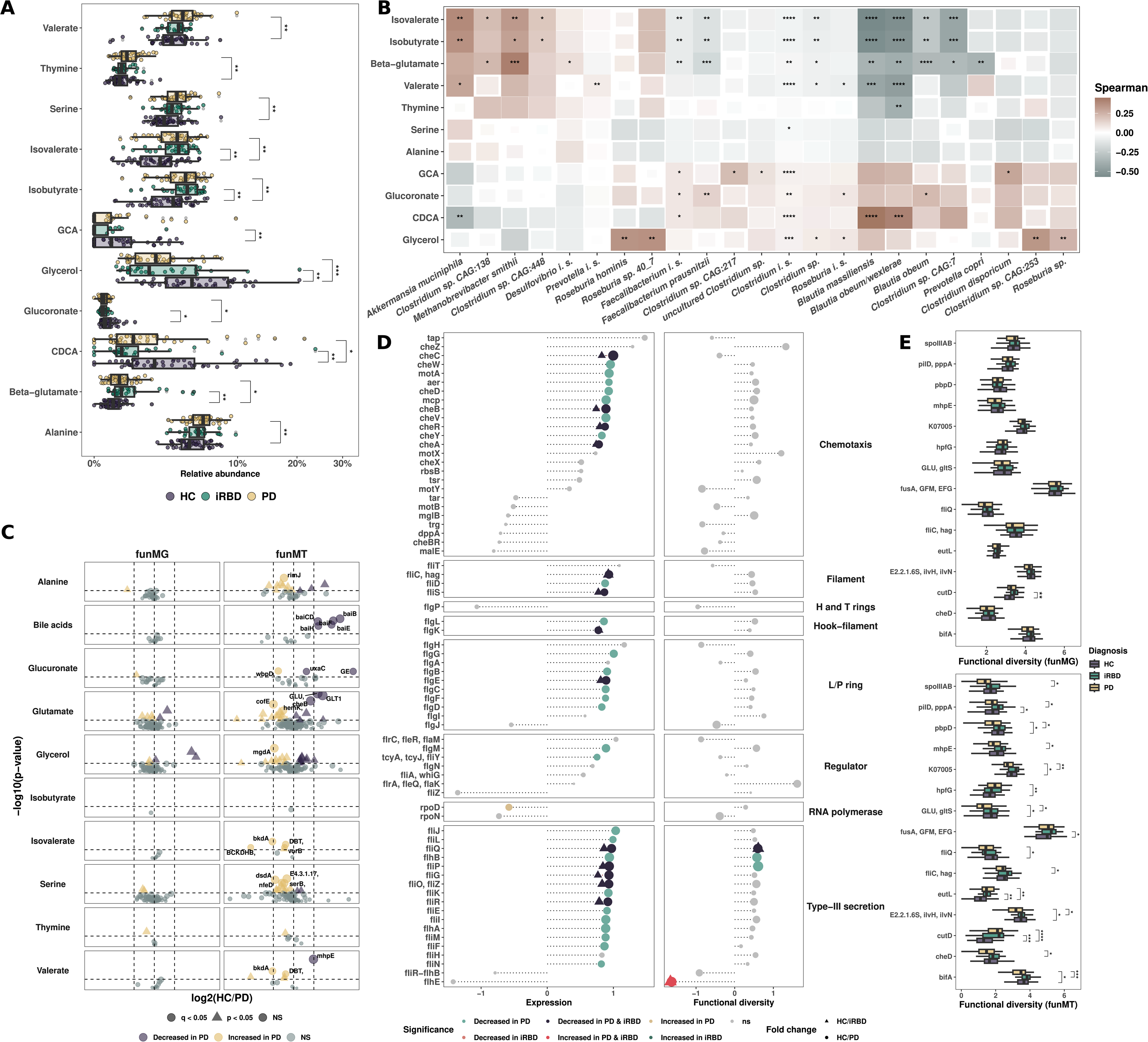
Altered metabolome is linked with microbial activity and transcripts. **A.** Metabolite relative abundances for significant compounds. Metabolomic data from untargeted meta-metabolomics, targeted SCFA and targeted bile acids were combined after normalization by sum for each. Dunn test, FDR corrected. **B**. Spearman correlation between taxMT at the species level with statistically significant metabolites. All p-values are FDR corrected. Genera are selected based on differential abundance or relevance in the literature. **C**. Absolute log2 fold change between HC and PD for funMG and funMT associated to significant compounds. Dots are scaled by the –log10(p-value), colorized and shaped according to p-value significance before (triangle shape) and after FDR correction (round shape). **D**. Chemotaxis and flagellin assembly pathway genes expression and Shannon index fold change between HC and PD group. Dots are colorized and shaped according to p-value significance before (triangle shape) and after FDR correction (round shape). Wilcoxon test, FDR corrected. **E**. Functional diversity comparison for funMT KO found as significantly different in pairwise differential analysis. Shannon index was calculated for each KO for funMG and funMT. Only genes significantly different after FDR correction on a Kruskal-Wallis test are plotted. P-values are calculated using Dunn post hoc test.

We next checked correlations between metabolite abundances and microbial abundances alongside activities. Metabolite abundances were linked to microbial abundances for taxMG and taxMT. TaxMG exhibited fewer significant correlations with metabolite abundances compared to taxMT, except for *Methanobrevibacter smithii* and *Clostridium* sp. CAG:273, which were positively correlated with β-glutamate levels (Extended fig. 3C, Spearman test, q < 0.01). Conversely, *Prevotella copri* and *Faecalibacterium prausnitzii* were negatively correlated with β-glutamate levels, while *Roseburia hominis* was positively correlated with glycerol levels (Extended fig. 3C, Spearman test, q < 0.001). Additionally, for taxMT we noted positive correlations with β-glutamate, isovalerate and isobutyrate with *A. muciniphila* and *Clostridium* sp*. CAG:448* alongside *Methanobrevibacter smithii* (Figure 3C, Spearman test, q < 0.01, q < 0.001 and q < 0.05, respectively), while *F. prausnitzii, Blautia massilliensis, Blautia obeum/wexlerae, Blautia obeum and Clostridim* spp. were negatively correlated with β-glutamate. In addition*, R. hominis, Roseburia* spp. and *Clostridium* spp. were positively correlated with glycerol (Figure 3C, Spearman test, q < 0.05). Of note, while performing the correlations at the genus level, we noted negative correlations between *Eubacterium* and isovalerate, isobutyrate and valerate (taxMT, q < 0.05, Spearman test, data not shown). Strikingly, *A. muciniphila*, some *Clostridium* spp. and *M. smithii* were positively correlated with compounds increased in PD while *R. hominis*, *F. prausnitzii, B.* species and *P. copri* were negatively correlated with these same compounds, highlighting groups of bacteria being linked with either individuals with PD and iRBD or HC (Figure 3C).

### Expression of genes linked to glutamate, bile acids and flagella is dysregulated in the iRBD and PD gut microbiome

To reinforce the results of the differential analysis, we acquired all the orthologs related to the metabolites which were statistically significant between the three groups. More specifically, we used regular expression matching to retrieve all orthologous genes that are linked to the above-mentioned metabolites in KEGG. Because the KEGG database only has one metabolite entry annotated as β-glutamate, we selected all glutamate-related genes instead (linked to both L-and D-glutamate). We found no statistically significant differences in gene abundances after correction between any of the three groups (funMG, Figure 3D and Extended fig. 3D, q > 0.05, Mann-Whitney test). However, we did find a decrease in transcripts in PD for the three known glutamate synthase genes (funMT, *GLT1*:K00264, *GLU*:K00284 and *gltB*:K00265, Figure 3D, Mann-Whitney test, q = 0.004, q = 0.004 and p = 0.006, respectively) alongside a decrease in cheB in PD (K03412, protein-glutamate methylesterase/glutaminase, Figure 3D, q < 0.01, Mann-Whitney test). Furthermore, we found an increase in cofE, mainly found in Archaea and involved in methanogenesis (K12234, coenzyme F420-0:L-glutamate ligase, q < 0.05, Mann-Whitney test). In addition, we found a decrease in BA-related transcripts in PD while transcripts related to serine and isovalerate were increased in PD (Figure 3D, q < 0.05). HC vs iRBD comparisons revealed significant differences in alanine-related transcripts but not for the other metabolites (Extended fig. 4A, q < 0.05). β-glutamate abundance was negatively correlated with the transcripts of the three glutamate synthases and carbamoyl-phosphate synthases, but positively correlated with methyl aspartate mutase, methylamine-glutamate N-methyltransferase and glutaminase (Extended fig. 4B, Spearman correlation test, q < 0.01).

Since *cheB* is part of the chemotaxis and flagellar assembly (FA) KEGG pathways, we further inspected these pathways to assess the microbial capacity for motility. We found no statistically significant differences in gene abundances (funMG) for either flagellar assembly or chemotaxis pathways between the three different groups (data not shown, q > 0.05). In contrast however, based on funMT, the chemotaxis pathway (14/26 transcripts decreased in PD, 0/26 transcripts increased, Figure 3E, q < 0.05) and the FA pathway (25/46 transcripts decreased in PD, 1/46 transcripts increased in PD, Mann-Whitney, Figure 3E, q < 0.05) were strongly downregulated in PD. In addition, we found transcripts decreased in both PD and iRBD for five transcripts belonging to chemotaxis and eight belonging to flagella assembly (q < 0.05, Mann-Whitney, Figure 3E). Moreover, eight transcripts showed a decrease in alpha diversity for chemotaxis and flagellin assembly pathways (p < 0.01, Mann-Whitney, Figure 3E). Finally, we observed a decrease in alpha diversity for *GLU*, flagellar assembly transcripts (*fliJ*, *fliQ* and *fliC*) and *cheD* in PD compared to the other two groups (Figure 3F, q < 0.05, Dunn test) while *pilD* and *aaaT* were elevated in iRBD (Figure 3F, Dunn test, p < 0.05).

### Flagellin and chemotaxis are differentially expressed depending on taxonomy and disease status

We next assessed the taxa expressing secondary bile acids biosynthesis (SBAS), FA and chemotaxis pathway genes. While resolving taxonomically SBAS at the genus level we found an interesting divergence between the differential analysis at the transcript the taxonomic level. Indeed, we previously found that *baiCD*, *baiE*, *baiB*, *baiH* and *baiF* were all significantly decreased in PD, while *baiA* and *baiN* were not (Figure 2C). We found 17 statistically significant differences in transcripts for *baiA* and *baiN* after FDR correction and 31 before correction when resolving their expression at the genus levels, but not for the other transcripts previously found as differentially expressed (q < 0.05 and p < 0.05, respectively, Extended fig. 5A). We found a significant decrease of expression for *baiN* in PD for *Blautia*, *Fusicatenibacter*, *Choladocola*, *Oliverpabstia* and *Ruminococcus* genera and an increase in expression for *baiA* encoded by *Lawsonibacter* genus in PD (q < 0.05, Mann-Whitney test, Extended fig. 5A).

When resolving the taxonomic expression of FA and chemotaxis genes we noted a differential expression according to disease and taxonomy. More specifically, we identified different clusters of taxa expressing FA and chemotaxis genes. The first cluster was composed of microbes expressing these genes principally in PD, including *Ruminiclostridium*, *Enterocloster*, *Dysosmobacter* and *Butyvibrio*. A second cluster was composed of microbes expressing these genes principally in HC, including *Roseburia*, *Agathobacter* and *Eubacterium* (Figure 4). Strikingly, we found that the third cluster is composed of taxa expressing flagellin or chemotaxis genes only in PD, including *Escherichia*, *Cellulosilyticum*, *Citrobacter* or *Eisenbergiella*. A fourth cluster was composed only of taxa expressing in HC including *Flavonifractor*, *Succinivibrio*, *Eisenbergiella* or *CAG-603* (Figure 4). We subsequently investigated the expression levels of genes of the extracellular parts of flagella in the cluster wherein *Roseburia* and most *Lachnospiraceae* were located (Cluster 2). Overall, we found a decrease in flagellin (*fliC*), filament cap (*fliD*, *fliS*) and hook-filament junction genes (*flgK*, *flgL*) in PD or iRBD for *Roseburia*, CAG-115 and *Agathobacter* genera (Extended fig. 5B, p < 0.05, Mann-Whitney test).

**Figure 4.**
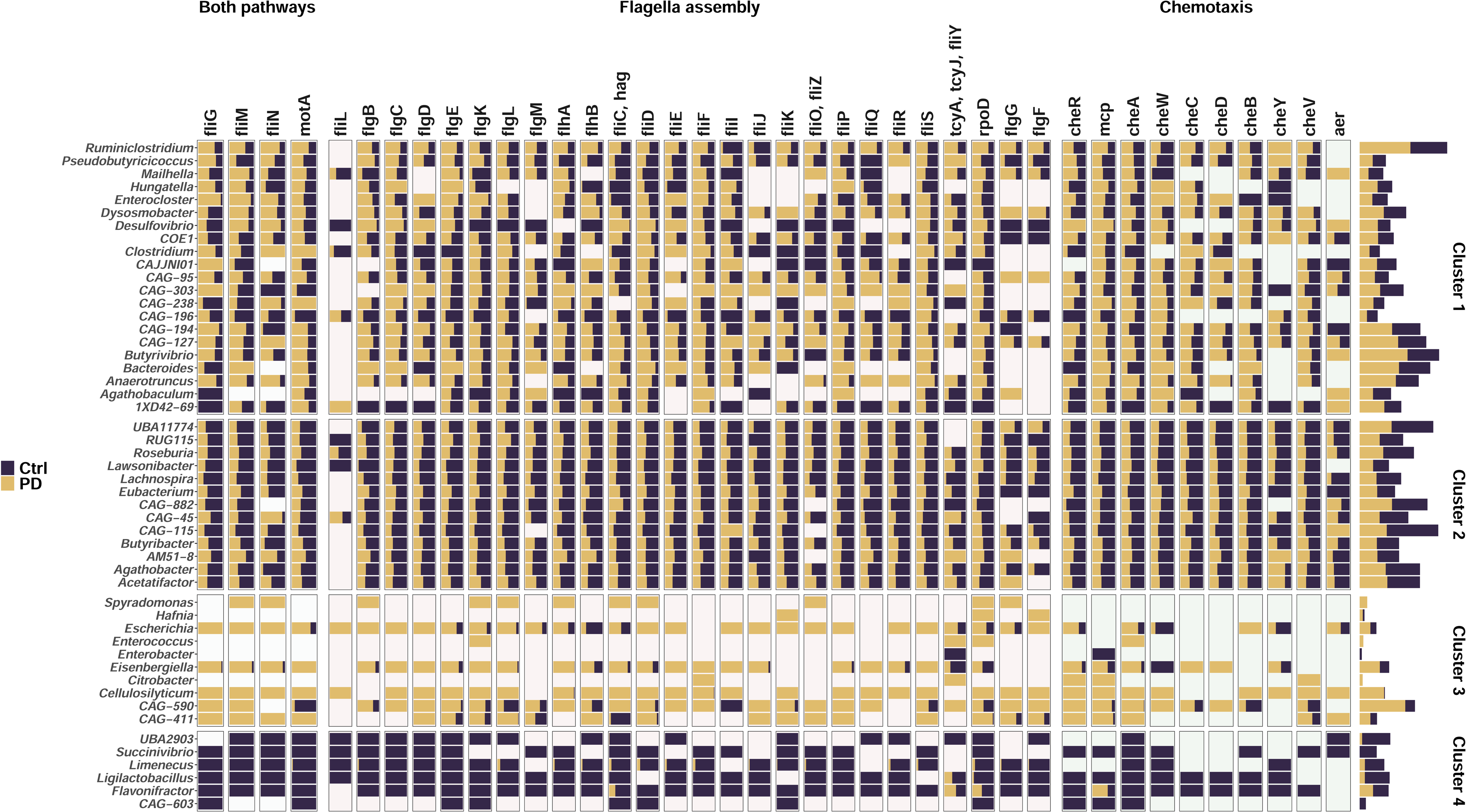
Flagellar assembly and chemotaxis gene expressions according to taxonomy. Top 50 genera expressing flagellar assembly and/or chemotaxis related genes are depicted by the mean of transcripts per million per disease status. Values are depicted for each genus and genes on the left panel and the sum of all genes per genus on the right panel with a square root transformation. Genera are clustered based on the log2FC(HC/PD) for each gene using a Canberra distance and *hclust()* using “ward.D2” method.

### Metabolites and metabolites-related genes associated with PD are central to the microbial ecosystem metabolism

To quantify the importance of glutamate derivatives and related genes in microbial metabolism, we next reconstructed microbiome-wide metabolic networks as previously described^34^. We mapped genes related to the compounds identified earlier as significantly different between the different groups. The metabolic network highlighted glutamate-related genes as compact and placed in the middle of the network while glycerol was more scattered across the network (Figure 5A). Strikingly, glutamate-related genes formed a subnetwork central to the overall network with a betweenness centrality measure (BC) of 93.0 compared to 0.0009 for the whole community (Figure 5B). Crucially, L-glutamate was the most central non-cofactor metabolite in the network and 2-oxoglutarate (L-glutamate derivative) was the second most central metabolite (Figure 5C). Considering this, it is apparent that glutamate and glutamate-related genes are central to microbiome metabolism and that modifications in the levels of these metabolites or transcripts reflect profound modifications of microbial metabolism.

**Figure 5.**
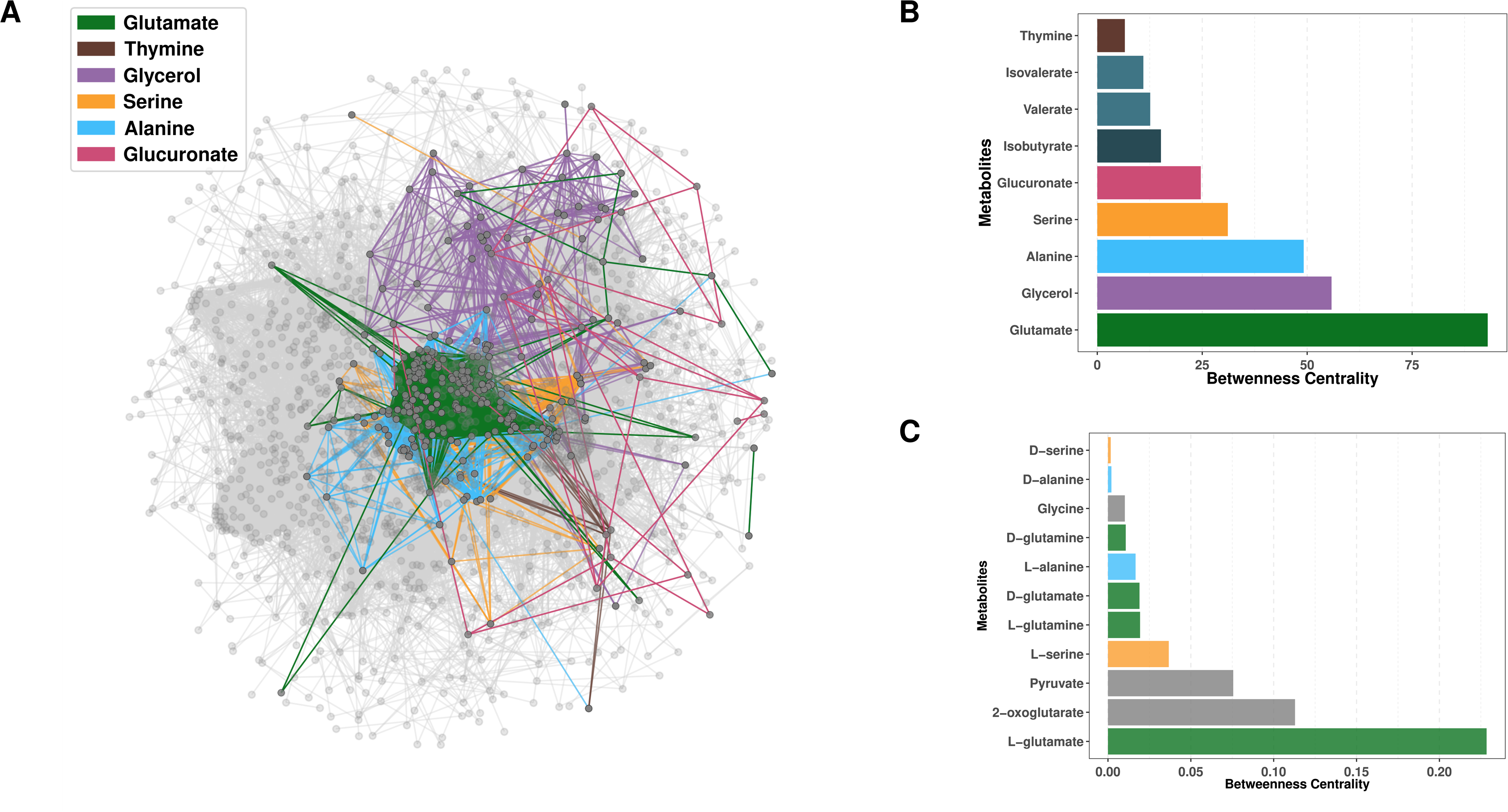
Metabolic network of whole community interactions. **A.** Metabolic network of whole community interactions, with KEGG KOs represented as nodes and associated metabolites as edges. Node sizes reflect MT/MG ratio of normalized read counts for each KO. Highlighted is the overlap between glutamate-, thymine-, glycerol-, serine-, alanine-and glucuronate-associated subnetworks mapped on the whole community network. **B.** Betweenness Centrality calculated for the key metabolites highlighted in the whole-community network based on genes as nodes. **C.** The network was inverted to calculate Betweenness centrality for metabolites, here metabolites are nodes and genes are edges.

### Multi-Omics Factor Analysis validates β-glutamate and flagella links with PD

To validate our findings, we used an unsupervised method with the Multi-Omics Factor Analysis (MOFA). The resulting MOFA model included 10 factors (F1-10 hereafter) whereby F1 showed a strong association with disease status (p < 0.05, ANOVA, Figure 6A). A complete description of the MOFA model is provided in the Extended information. We found that F1 mainly explained the variance of funMT, taxMT, and MM (17.6%, 9.6%, and 5.7%, respectively, Extended fig. 6A-B) and showed separation of HC and PD, but not iRBD versus other groups (p < 0.05, ANOVA, Figure 6B). Specifically, the microbiome of PD was characterized by the joint increase in abundance of *M. smithii*, archaeal proteins and genes (based on MP and MG data), *A. muciniphila* (based on taxMT data), β-glutamate, isovalerate, isobutyrate, hexadecanoic, and hyocholic acids, whereas the abundance of *R. hominis* (taxMG data), flagellin (funMT), GCA and glycerol were decreased in PD (Figure 6C). Overall, the MOFA results are strongly consistent with the per omic layer analysis and validate these findings.

**Figure 6.**
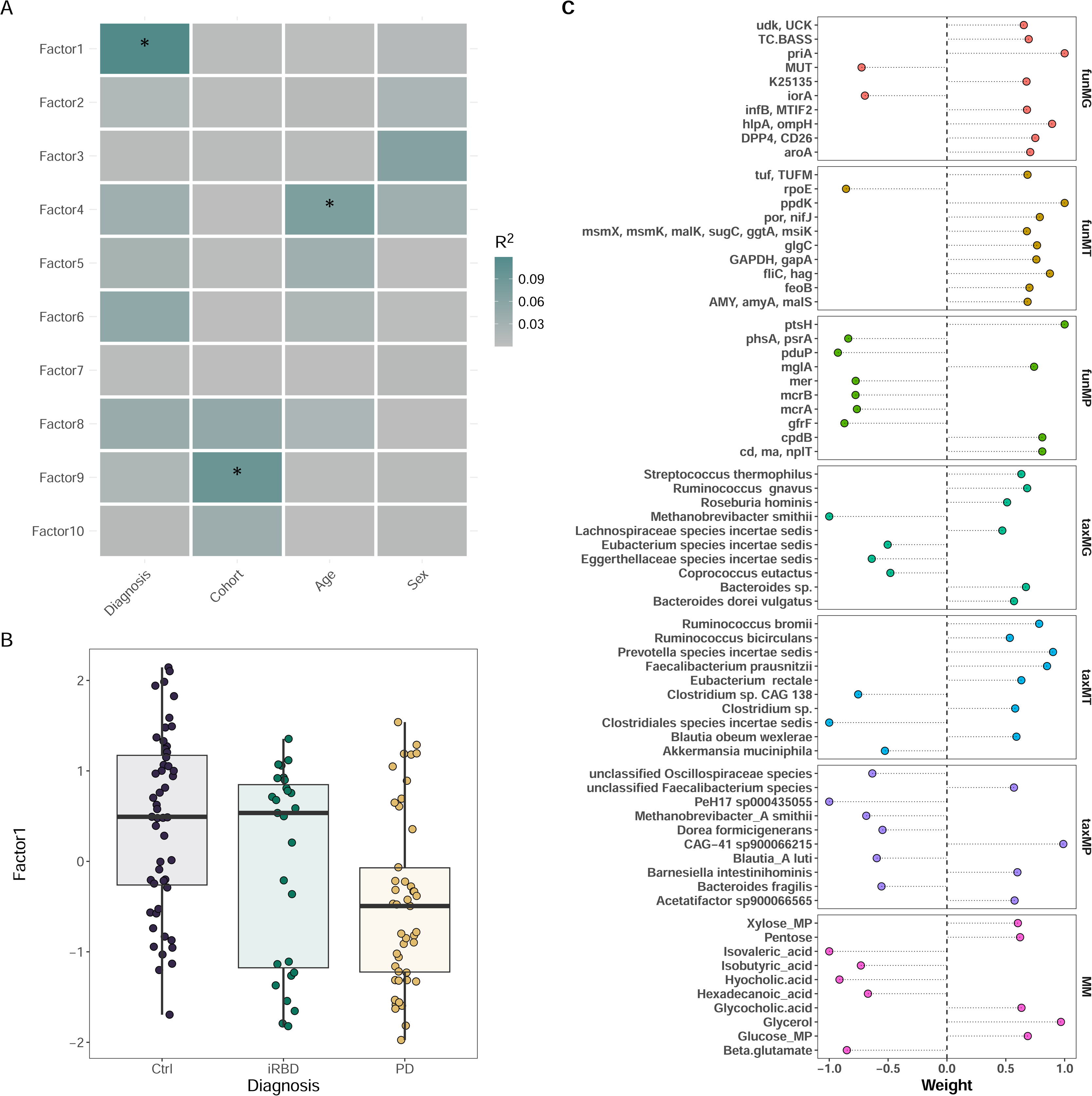
MOFA analysis validates the findings of per omics layer analyses. **A.** Associations of MOFA factors with the diagnosis and confounders, the color of rectangles represents partial R² values, significant associations (FDR-adjusted p-value < 0.05) marked with an asterisk. **B.** Abundance of the Factor 1 in studied groups. **C**. Min-max scaled weights of top 10 features per omics layer contributing to Factor 1. The sign of the weight indicates the direction of the effect, the abundance of features with positive weights is positively associated with the Factor 1 level, and the abundance of features with negative weights is negatively associated with the Factor 1 level.

### From the microbiome to the host: expanding the Expobiome map

To contextualize our newly gained insights about the links between microbial taxa, their molecules, and mechanistic evidence from the literature on host physiology, we integrated our results into the Expobiome map^35^ (https://expobiome.lcsb.uni.lu). Specifically, we added *R. intestinalis* and *hominis*, *B. wexlerae, coccoides* and *producta*, *Clostridium butyricum* and *F. plautii* as new taxonomic elements linking flagellin, lipoteichoic acid (LTA) and SCFAs signaling to immune-relevant human proteins (Figure 7). We also added mechanistic links on *F. prausnitzii* and its effects mediated by the short-chain fatty acid butyrate. Thereby, we integrated new mechanistic details linking the combinatorial functions from these taxa and pro-inflammatory components effectors such as TLR5 and its downstream activation of NLRP3 inflammasome and NF-κB, IL-6 and TNF-α secretion. Thereby these new additions to the Expobiome map highlight the convergent immunomodulatory functions of commensal taxa, perturbed in the context of PD.

**Figure 7.**
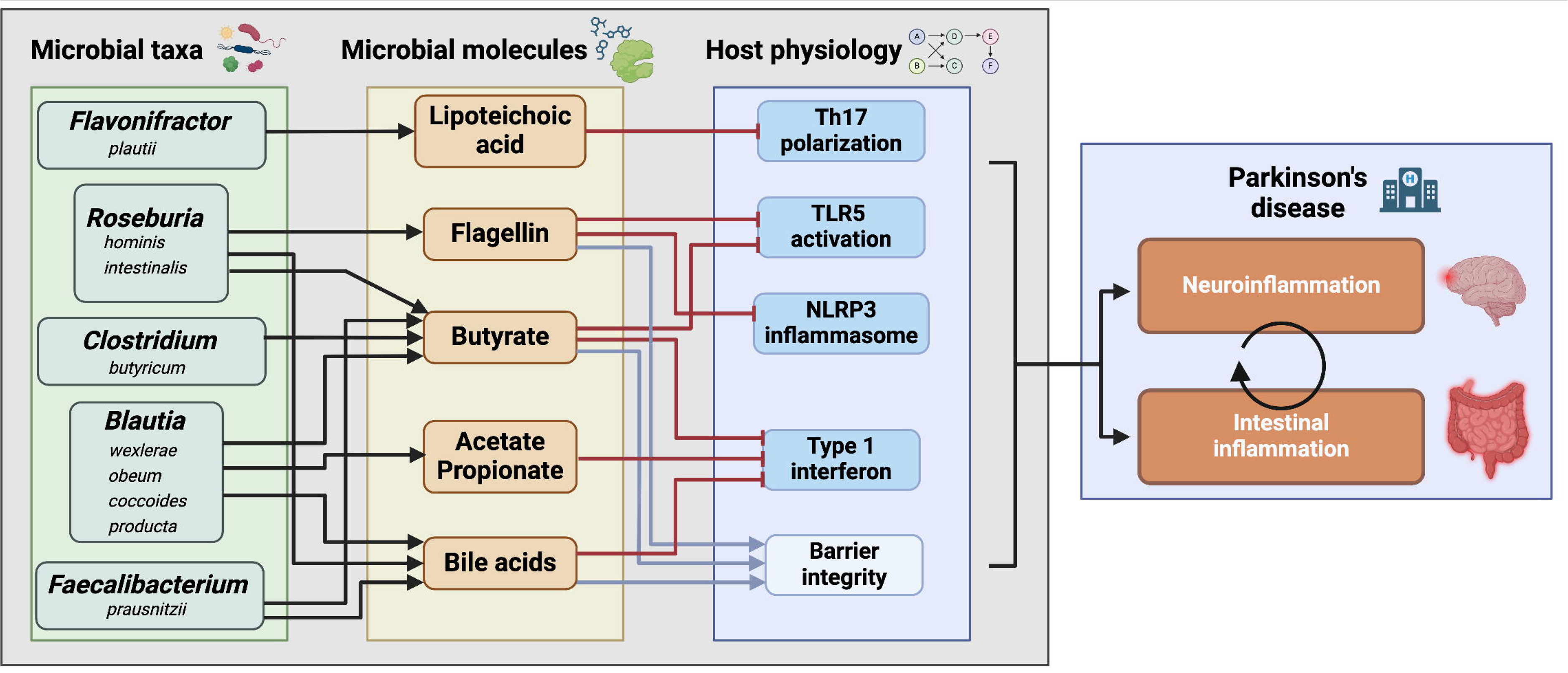
Schematic representation of the Expobiome map new taxonomic and molecular additions. Schematic representation of the new additions to the Expobiome map. In short, we collected literature highlighting mechanistic evidences linking microbes to the host physiology. We integrated *Roseburia*, *Blautia*, *Faecalibacterium*, *Flavonifractor* and *Clostridium* species as new taxonomic elements and linked their molecules to the host physiology. Finally, the Expobiome map is now connected to Parkinson’s disease map, providing direct overlay of the shared proteins between the maps.

To facilitate the investigation of potential relationships between microbial taxa, microbial compounds, and human proteins that have been identified as crucial players in PD, we have furthermore linked the Expobiome map to the Parkinson’s disease map (https://pdmap.uni.lu). Notable overlaps exist between host proteins that have been established to interact with microbial components in both the immune system and the central nervous system. For example, dendritic cells and macrophages have well characterized interactions with flagellin through TLR5 binding. TLR5 and other antigen sensing receptors are present on microglia, a cell type of critical importance in neuroinflammation and PD. This integration of information from both thus-far disconnected maps allows for a more comprehensive understanding of these interactions and helps to draw new hypotheses on the intricate links between the gut microbiome, the immune system and PD.

## Discussion

Here we investigate the links between the gut microbiome and PD using an integrated multi-omics approach based on standardized sample collection and extraction. We include individuals with iRBD as a prodrome of PD to compare early and later stages of the disease but do not find statistically significant differences between iRBD and PD, especially in comparison to the more pronounced differences found between HC and PD. Previous studies have shown differences in the different early stages of PD, but these studies were performed using 16S rRNA gene amplicon data^7,10^. In contrast to the amplicon-based results, no differentiation between iRBD and PD are found using metagenomic data^5^. More specifically, and contrary to previous findings^5^ including our own work^7^, we do not find significant differences between HC and PD individuals in the metagenome with respect to alpha or beta diversity. However, we show differences in transcriptional activity for *R. hominis*, *B. obeum, B. wexlerae* and *P. copri* which are known to be found at lower abundance in PD^5,6,8–13^.

Moreover, we find several species, including *A. muciniphila*, *M. smithii* and *Clostridium* spp, to be positively correlated with the abundance of compounds increased in PD such as β-glutamate, isovalerate or isobutyrate, while other species are inversely correlated with these compounds (species belonging to *Faecalibacterium*, *Blautia*, *Eubacterium* and *Roseburia*). Collectively, these findings are in line with findings associating those genera with PD^5,6,8–13^.

We identify metabolites that are associated with PD, amongst which BAs, alanine, serine and β-glutamate showed differences in linked gene expression. The BA chenodeoxycholic acid (CDCA) is decreased in PD and iRBD whereas glycocholic acid (GCA) is decreased only in iRBD. We also detect a decrease in transcripts related to BAs in PD but not iRBD. BAs have a wide spectrum of effects on the immune system^22,23^, metabolism^35^, hormones^36^ and on the CNS^37^. GCA has been found to be increased in the CNS of PD and associated with disease duration^37^, stressing the importance of analyzing BAs in PD and prodromal stages of PD. Interestingly, when we taxonomically resolve the differentially expressed SBAS genes, we find different signals than when we compare the genes by themselves. Indeed, for the differential expression analysis we found a significant decrease in PD for *baiB*, *baiCD*, *baiE*, *baiF* and *baiH* but not for *baiA* and *baiN*, when we resolved the taxonomic expression of the SBAS genes we find differences only for *baiA* and *baiN*. This finding underscores the importance of considering the microbiome at the taxonomic and functional level at once. In contrast to previous studies^6,15^, we do not find a decrease in butyrate, propionate, acetate or overall levels of SCFAs. However, we find an increase in isovalerate, isobutyrate and valerate as described previously^15^. Previous studies have highlighted the correlation of fecal concentrations of isobutyrate and isovalerate with PD severity^38^ and concentrations of valerate with disease duration^39^. There currently is an apparent lack of knowledge regarding isobutyrate or isovalerate with respect to host-microbiome interactions and very few related genes annotated in the KEGG database, thereby hindering interpretations concerning potential links between these metabolites and PD.

β-glutamate and glutamate-related genes are of particular interest in the context of PD because of the reported toxic effects of L-glutamate on neurons^24^ and because of its association with microbial activity. In addition, glutamate levels have been reported to be increased in PD individuals’ blood sera^17,18^. β-glutamate has strikingly only one reaction described in the KEGG database, in contrast, L-glutamate has 213 reactions and 54 pathways, while D-glutamate has 12 reactions and 2 pathways. We find that glutamate and glutamate-related genes are central to microbial metabolism, which underpins the notion that the highlighted differences reflect a pronounced impact on gut microbiome function. Of note, glutamate likely has local effects on enteric neurons with subsequent influences on the CNS^40^. Although we show significant differences in β-glutamate and glutamate-related genes in the stool of PD patients, we detect no significant differences in the glutamate levels (both enantiomers) between the three groups. We highlight significant differences in β-glutamate for which we currently cannot evaluate the effects on the ENS and CNS. Overall, based on our results, microbiome-driven glutamate metabolism and its impact on the glutamatergic system must be comprehensively studied in the future to disentangle its link with PD.

Glutamate-related genes are further involved in chemotaxis and flagellar assembly pathways, highlighting a modification of the latter genes’ expression with the majority being decreased in PD individuals and some of these genes in iRBD too. A decrease in flagellar assembly gene abundances has been previously reported in a metagenomic analyses of PD^9^. We do not see significant differences in the linked gene abundances, but their expression levels are significantly different, highlighting altered regulation of transcription in PD/iRBD compared to HC. Flagellar assembly and chemotaxis genes are also differentially expressed by specific microbes; the genera *Escherichia* and *Cellulosilyticum* for instance are expressing flagellar assembly genes only in the context of PD without being statistically differentially abundant.

Flagellin is a known immunogenic molecule, a potent pro-inflammatory compound in pathogens^41–43^ and is thus targeted by secretory IgA^44^. However, flagellin in commensals, especially in the *Lachnospiraceae* family, has been shown to be either ‘silently recognised’^45^ or elicit anti-inflammatory effects^46–48^. Amongst the *Lachnospiraceae* family, the genus *Roseburia* shows a decrease in the transcription of flagellin in the gut microbiome of PD. This in turn may be linked to immune system dysregulation and exert indirect effect on the CNS, particularly in microglia as shown in a previous study using a murine model of PD^49^. Previous studies have shown an increased inflammatory state in PD, with increased pro-inflammatory circulating immune cells^50^, cytokines^51^, and activated microglia^52^. In addition, microglia can be activated by α-synuclein via NLRP3 following TLR2 (lipopolysaccharide (LPS) sensing) and TLR5 (flagellin sensing) activation^53^. Therefore, the activation or inhibition of TLRs by distinct flagellins may modulate microglia activation by competing with α-synuclein and affect PD progression. It is however not known if flagellin can reach the CNS directly or if this potential effect would be mediated by an indirect effect through the immune system. Evidence exists that a local, microbiome-mediated effect on the enteric immune system can impact distant sites such as the brain^54^ or tumors^55^. This provides solid grounds for the hypothesis that flagellin can regulate the CNS and modulate the neuroinflammation occurring in PD as resolved in the Expobiome map (Figure 7).

Knowledge on microbial antigens is based on studies of pathogens, mainly *Salmonella typhimurium*, pathogenic *Escherichia coli*, *Staphylococcus aureus* or *Bacillus subtilis*. The results of these studies have led the scientific community to consider bacterial antigens as pro-inflammatory and danger signals for the host^56^. Our findings associating flagellin expression with health, along with recent reports showing the anti-inflammatory effects of flagellin from *R. hominis* and *intestinalis*^45,46^, LTA from *F. plautii*^57^ and LPS from *Bacteroides dorei*^58^, call for a re-evaluation of the impact of common antigens on host physiology. Finally, the diverse compounds produced by the gut microbiome demand detailed investigation, with regard to the taxonomy and commensal/pathogen phenotype of the producing microbe, to fully understand their modulating effects on the immune and nervous systems alongside metabolism in health and disease.

In conclusion, our work clearly highlights the importance of studying microbiome functions rather than restricting microbiome analyses to taxonomic structure. Specifically, the combination of MT and MM provides clear insights into the activity of specific microbial taxa in relation to disease. In our present work, MT reveals that disease association is not solely determined by gene expression levels; the diversity of microbes capable of expressing a specific function and the specific taxa expressing those functions are also of immediate interest and relevance. While microbiome interventions are mainly designed to deal with dysbiosis at the taxonomic level, we propose here to focus on restoring the lost functions from dysbiotic gut microbiomes. The future of microbiome research might lie in understanding how we can modulate, re-activate or shut down specific microbial functions *in vivo* in order to improve knowledge and later improve or functionally tailor microbiome interventions.

## STAR Methods

### Key resources table

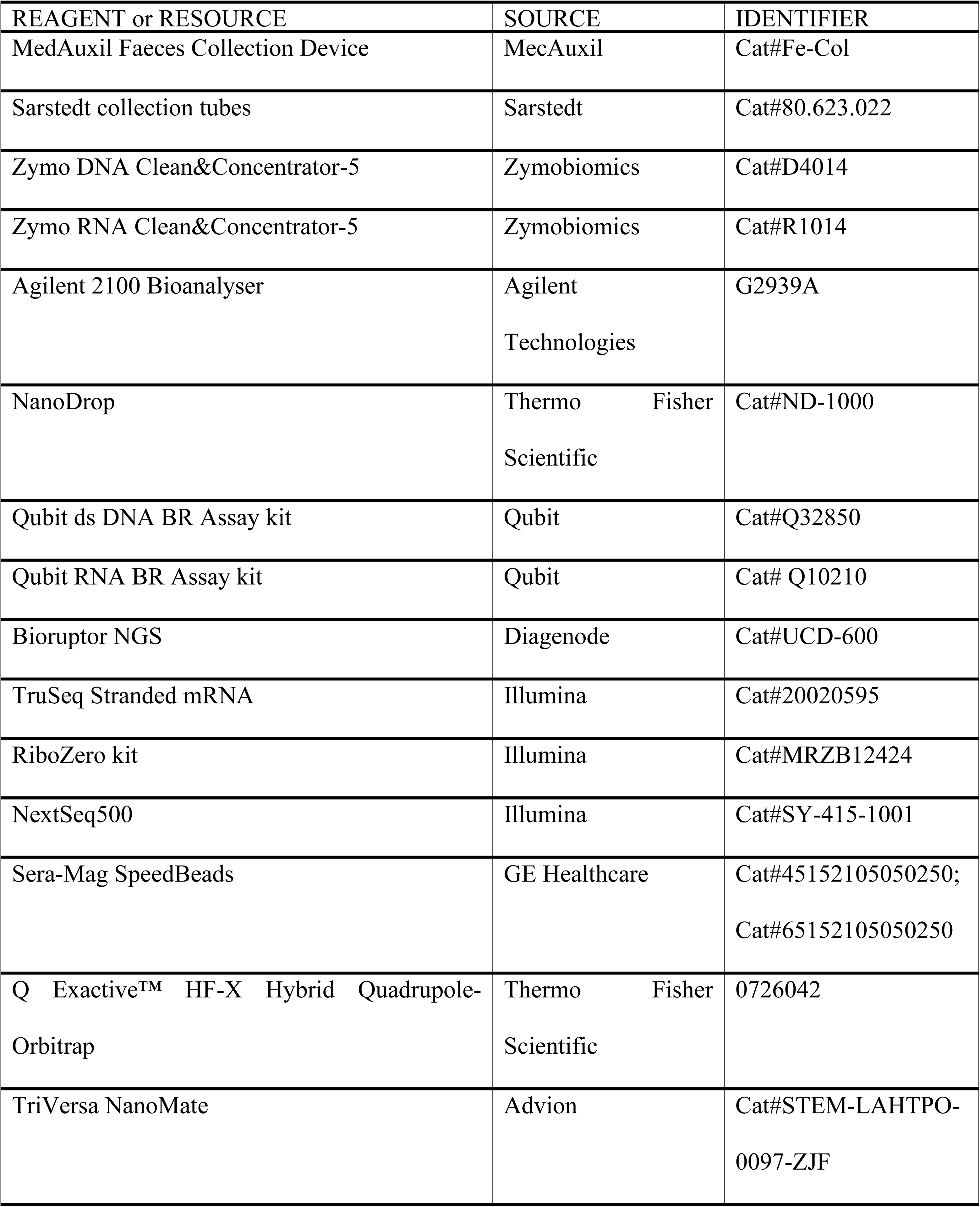
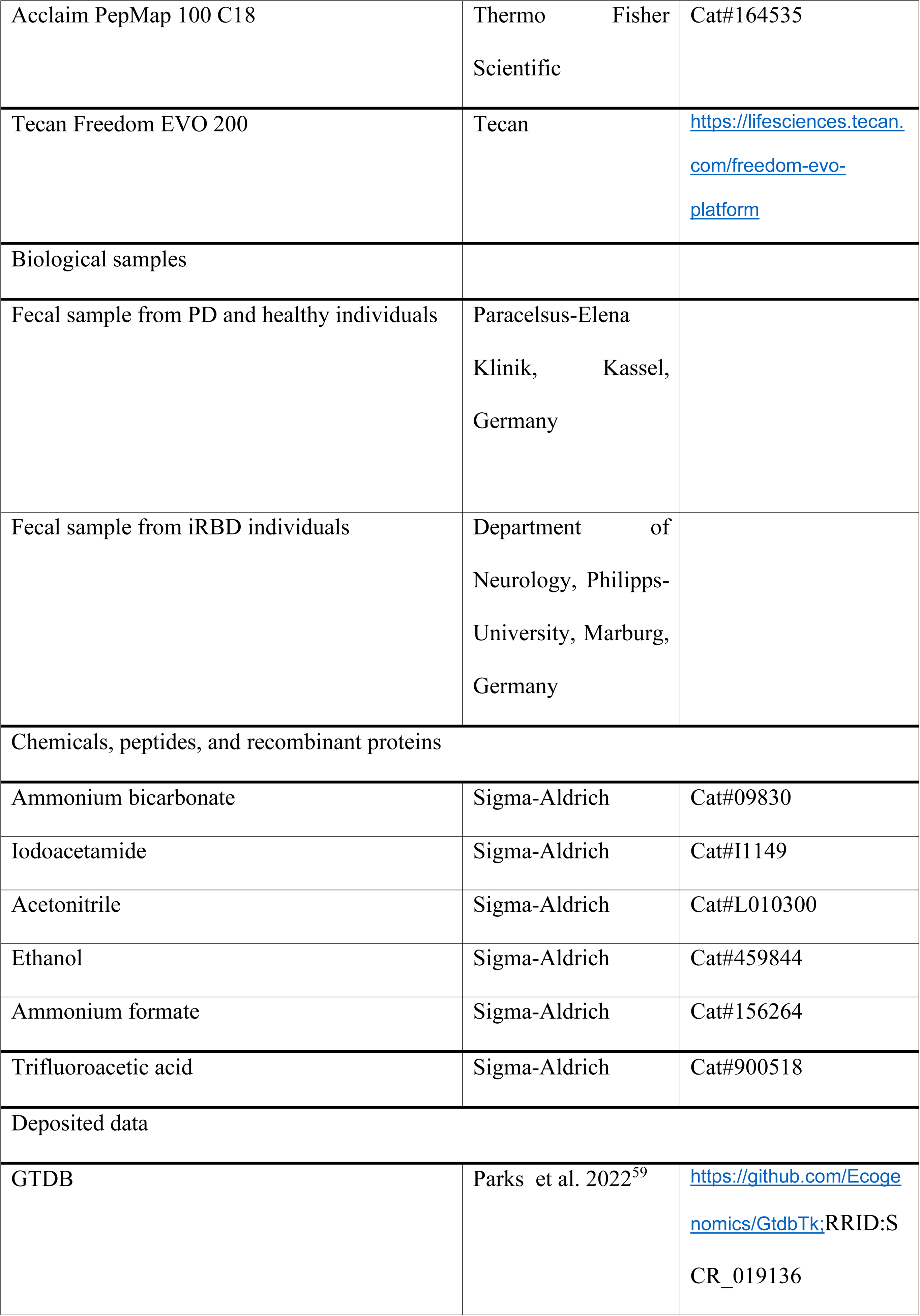
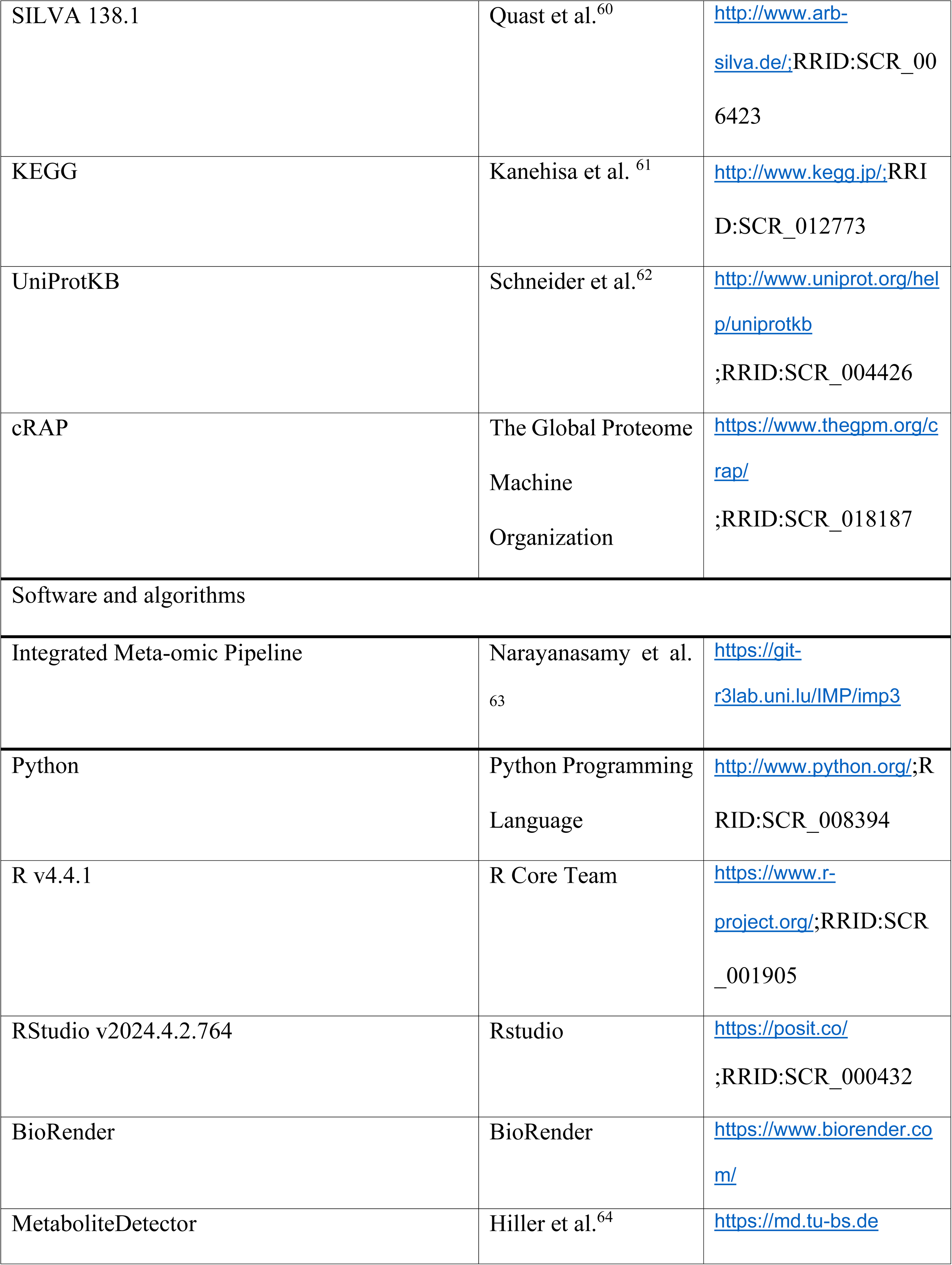
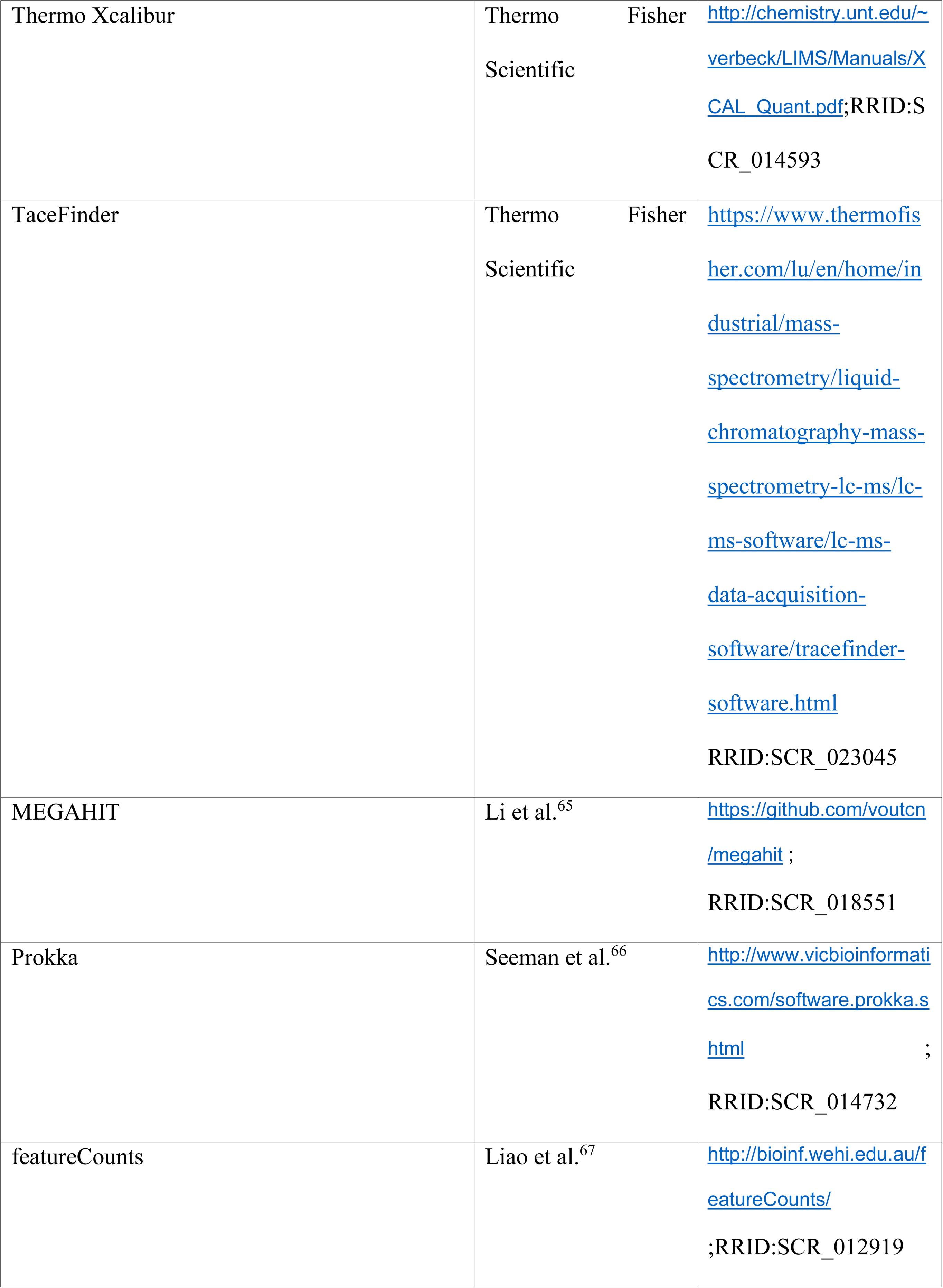
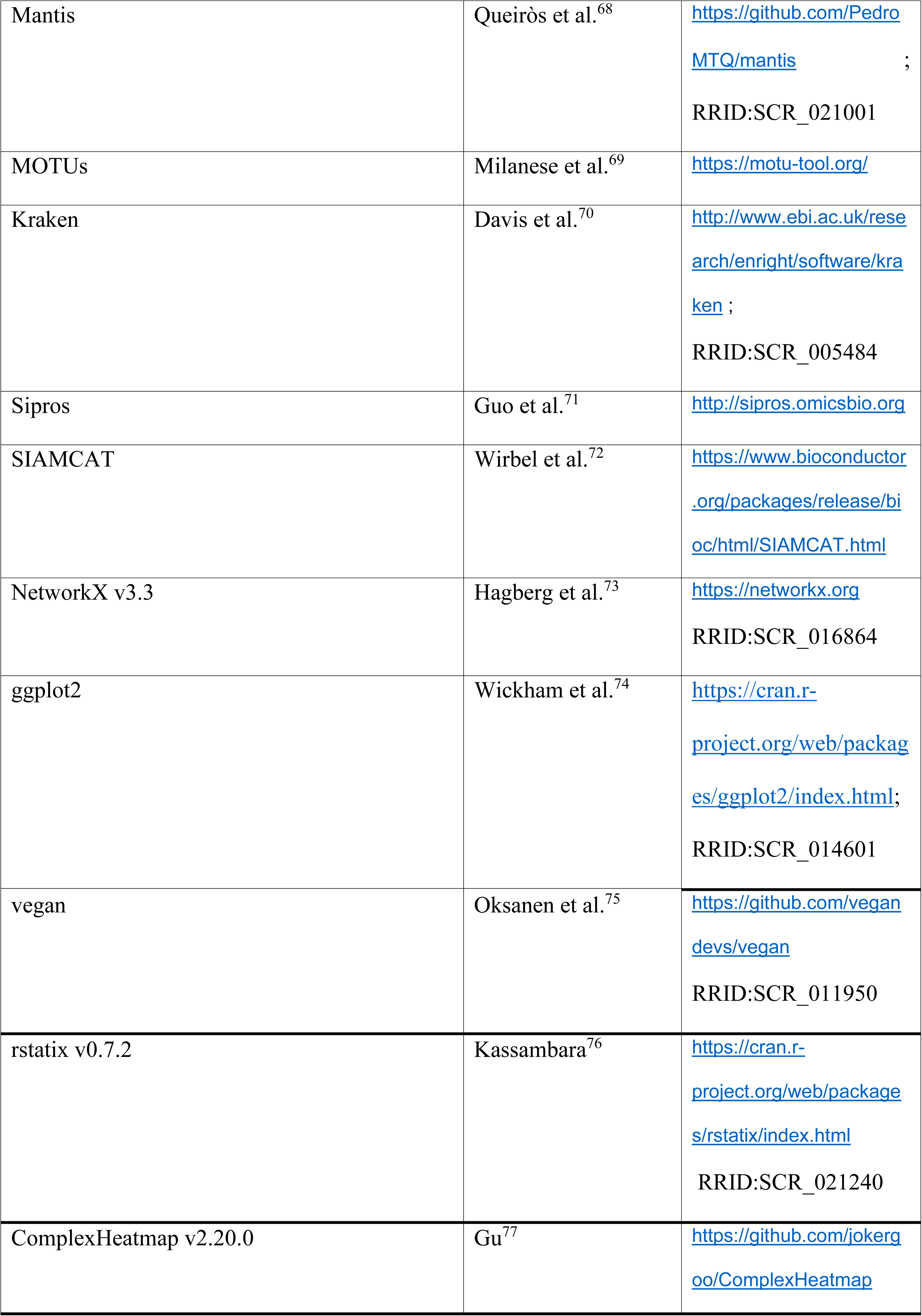
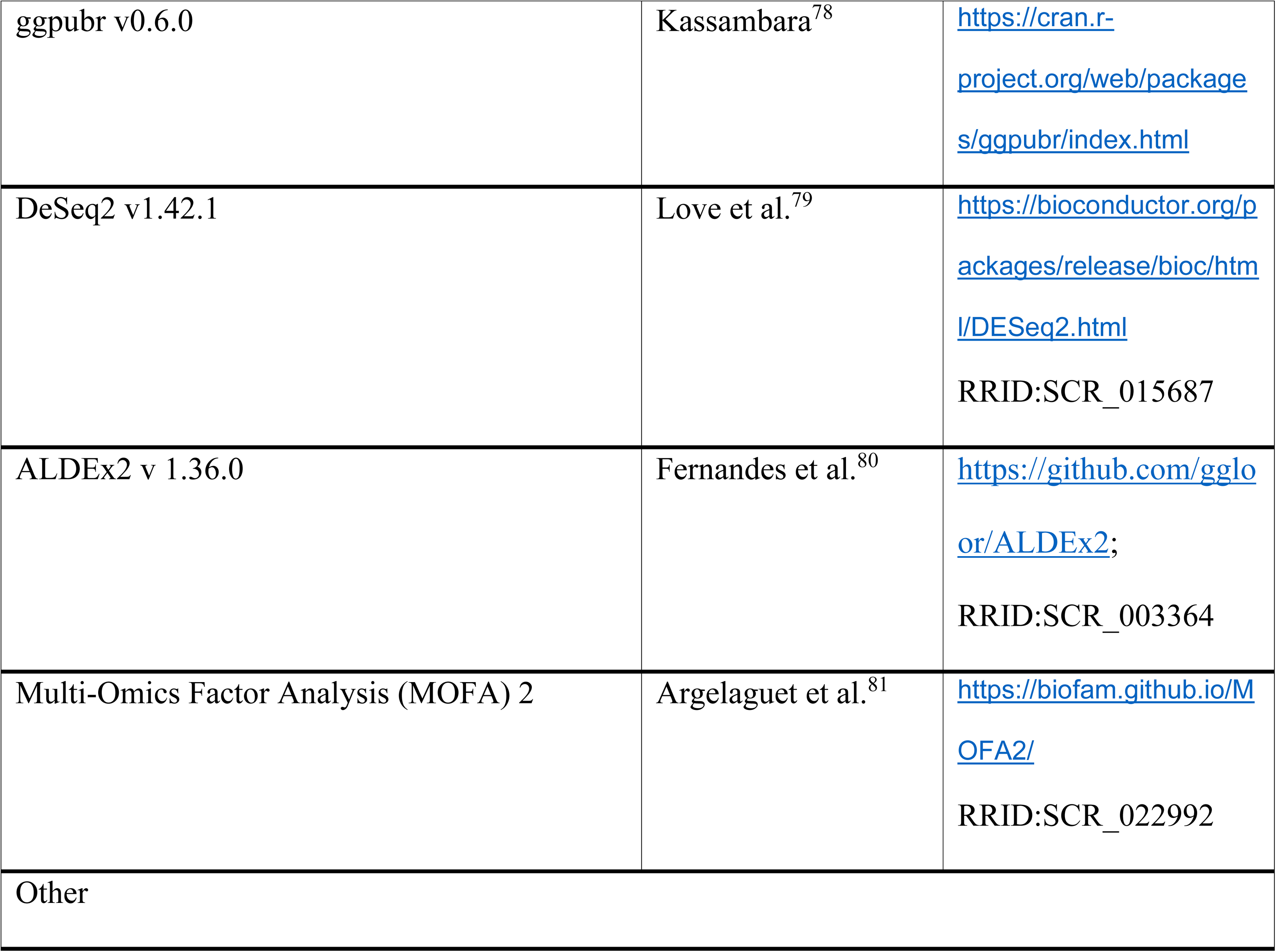

## EXPERIMENTAL MODEL AND STUDY PARTICIPANT DETAILS

All subjects from both cohorts provided informed written consent, and the sample analysis was approved by the Comité National d’Ethique de Recherche of Luxembourg (reference no.: 140174_ND).

### Kassel Cohort

The DeNoPa cohort represents a prospective, biannual follow-up study of (initially *de novo)* Parkinson’s disease (PD) patients at the Paracelsus-Elena Klinik, Kassel, Germany. Fecal samples from PD patients (46) and healthy controls (29) were collected during the 4-year follow-up visit for the cohort. Details on inclusion and exclusion criteria and ancillary investigations have been published previously^82,83^. Subjects with idiopathic rapid-eye-movement sleep behaviour disorder (iRBD, 13) were recruited at the same clinic, diagnosed according to consensus criteria of the International RBD study group^84^ using video-assisted polysomnography, and were included only if they showed no signs of a neurodegenerative disorder. DeNoPa subjects were required to have a 4-week antibiotic free interval before fecal sample collection. As additional control subjects, we collected fecal samples from (20) neurologically healthy subjects living in the same household as the DeNoPa participants. Samples of *de novo* PD patients from a cross-sectional cohort at the same clinic were included if subjects were recently diagnosed, drug-naïve and met United Kingdom Parkinson’s Disease Society Brain Bank (UKPDSBB) clinical diagnostic criteria^30^. All subjects except household HC were interviewed and examined by an expert in movement disorders. The study conformed to the Declaration of Helsinki and was approved by the ethics committee of the Physician’s Board Hessen, Germany (FF 89/2008). The DeNoPa trial is registered at the German Register for Clinical trials (DRKS00000540).

### Marburg Cohort

We also added samples from 14 patients with polysomnography-confirmed iRBD which were recruited from the outpatient clinic of the Department of Neurology, Philipps-University, Marburg, Germany, between November 2015 and November 2016. iRBD was diagnosed according to the guidelines of the American Academy of Sleep Medicine (AASM ICSD-3)^85^. A detailed medical history was recorded, and a complete neurological examination performed to verify the subjects’ suitability. Inclusion criteria were age above 18 years, no dopamimetic therapy, and no diagnosis of PD, MSA, DLB or PSP. Exclusion criteria were smoking, antibiotic therapy in the last 24 months, history of other neurological diseases or disorders of the gastrointestinal tract. Non-motor and autonomic symptoms were evaluated with the SCOPA-AUT^86^ and PD-NMS^87^ questionnaires. Motor function was evaluated with the UPDRS^88^. Additionally, patients were asked to complete the RBD-Sleep questionnaire^89^. The study conformed to the Declaration of Helsinki and was approved by the ethics committee of the Medical Faculty of the Philipps-University, Marburg, Germany (46/14).

## METHOD DETAILS

### Fecal sample collection

Fecal samples were collected at the clinics via a Faeces Collection Device (MedAuxil) and collection tubes (Sarstedt), as previously described^7^. Samples were immediately flash-frozen on dry ice after collection. Samples were subsequently stored at –80 °C and shipped on dry ice.

### Sample exclusions

The initial set of samples consisted of 50 PD, 30 iRBD and 50 healthy control subjects (HC). Three PD and two iRBD cases were subsequently excluded for clinical reasons (adjusted diagnosis), one iRBD and one PD subject for logistical reasons, and one control due to a combination of microbiome-altering medications (metformin, antidepressants, statins, and proton pump inhibitors). Additional samples were excluded due to missing values (metabolomics) or a low amount of identified analytes (metaproteomics), leading to the final numbers of samples summarized below:

‒ Metagenomics (MG) & metatranscriptomics (MT): 49 HC, 27 iRBD, 46 PD
‒ Metaproteomics (MP): 42 HC, 22 iRBD, 40 PD
‒ Meta-metabolomics: 49 HC, 27 iRBD, 41 PD

### Metagenomic and metatranscriptomic sequencing

Extractions from fecal samples were performed according to a previously published protocol^90^, conducted on a customized robotic system (Tecan Freedom EVO 200). After extraction, DNA and RNA were purified prior the sequencing analysis by using the following commercial kits respectively: Zymo DNA Clean&Concentrator-5 (D4014) and Zymo RNA Clean&Concentrator-5 (R1014). RNA quality was assessed and quantified with an Agilent 2100 Bioanalyser (Agilent Technologies) and the Agilent RNA 6000 Nano kit, and genomic DNA and RNA fractions with a NanoDrop Spectrophotometer 1000 (Thermo Scientific) as well as commercial kits from Qubit (Qubit ds DNA BR Assay kit, Q32850; Qubit RNA BR Assay kit, Q10210). All DNA samples were subjected to random shotgun sequencing. Following DNA isolation, 200-300 ng of DNA was sheared using a Bioruptor NGS (Diagenode) with 30s ON and 30s OFF for 20 cycles. Sequencing libraries were prepared using the TruSeq Nano DNA library preparation kit (Illumina) following the manufacturer’s protocol, with 350 bp average insert size. For MT, 1 µg of isolated RNA was rRNA-depleted using the RiboZero kit (Illumina, MRZB12424). Library preparation was performed using the TruSeq Stranded mRNA library preparation kit (Illumina) following the manufacturer’s protocol, apart from omitting the initial steps for mRNA pull down. MG and MT analyses, the qualities of the libraries were checked using a Bioanalyzer (Agilent) and quantified using Qubit (Invitrogen). Libraries were sequenced on an Illumina NextSeq500 instrument with 2×150 bp read length.

### Metaproteomics

20 µL protein extract were processed using the paramagnetic bead approach with SP3 carboxylate coated beads^91,92^. Briefly, the protein samples were reduced with 2µL 25 mM DTT in 20 mM ammonium bicarbonate (Sigma-Aldrich) for 1 h at 60°C. Subsequently, 4 μL 100 mM iodoacetamide (Merck) in 20 mM ammonium bicarbonate was added and incubated for 30 min at 37°C in the dark. Next, 5 μL of 10% formic acid was added as well as 70 μL 100% acetonitrile (ACN) to reach a final organic content higher than 50% (v/v). 2 μL SP3 beads per sample were washed with water three times with subsequent addition of the sample. After protein binding to the beads, the supernatant was discarded. The beads were washed twice with 200 μL 70% (v/v) ethanol, and once with 200 µL ACN. The protein lysates were proteolytically cleaved using trypsin (1:50) over night at 37 °C. Since trypsin is added in aqueous solution to the samples, the proteins are not bound to the beads during enzymatic cleavage. ACN was added to each sample to reach a final organic content higher than 95% (v/v). After peptide binding to the beads, the samples were washed with pure ACN on the magnetic rack. Finally, the peptides were eluted in two steps. First, with 200 μL 87% ACN (v/v) containing 10 mM ammonium formate (pH 10), and next with two times adding 50 μL water containing 2 % (v/v) DMSO and combination of the two aqueous supernatants. Thus, two fractions of peptides were generated, which were evaporated and re-dissolved in water containing 0.1 % formic acid (20 µL) and analyzed on a Q Exactive HF instrument (Thermo Fisher Scientific) equipped with a TriVersa NanoMate source (Advion) in LC chip coupling mode. Peptide lysates were injected on a trapping column (Acclaim PepMap 100 C18, 3 μm, nanoViper, 75 μm x 2 cm, Thermo Fisher Scientific) with 5 μL/min by using 98% water/2% ACN 0.5% trifluoroacetic acid, and separated on an analytical column (Acclaim PepMap 100 C18, 3 μm, nanoViper, 75 μm x 25 cm, Thermo Fisher Scientific) with a flow rate of 300 nL/min. Mobile phase was 0.1% formic acid in water (A) and 80 % ACN/0.08 % formic acid in water (B). Full MS spectra (350–1,550 *m/z*) were acquired in the Orbitrap at a resolution of 120,000 with automatic gain control (AGC) target value of 3×10^6^ ions.

### Meta-metabolomics

Untargeted GC-MS as well as targeted measurements (SCFA GC-MS/MS and bile acids LC-MS/MS) from fecal samples were performed according to a previously published protocol^93^. All GC-MS chromatograms were processed using MetaboliteDetector, v3.220190704^94^ while LC-MS chromatogram were acquired with Thermo Xcalibur software (version 4.1.31.9) and analyzed with TaceFinder (Version 4.1). Compounds were initially annotated by retention time and mass spectrum using an in-house mass spectral library. Internal standards were added at the same concentration to every medium sample to correct for uncontrolled sample losses and analyte degradation during metabolite extraction. The data was normalized by using the response ratio of the integrated peak area of the analyte and the integrated peak area of the internal standard.

### Sequencing data processing and analysis

For all samples, MG and MT sequencing data were processed and hybrid-assembled using the Integrated Meta-omic Pipeline (IMP)^95^ (https://git-r3lab.uni.lu/IMP/imp3, commit 8c1bd6fa443d064511909c9eede20703f45e6c69). It includes steps for the trimming and quality filtering of the reads, the filtering of rRNA from the MT data, and the removal of human reads after mapping against the human genome (hg38). Pre-processed MG and MT reads were assembled using the IMP-based iterative hybrid-assembly pipeline using MEGAHIT^96^ 1.0.3. After assembly, the prediction and annotation of structural features such as open-reading frames (ORFs) was performed using a modified version of Prokka^66^ and followed by functional annotation of those using Mantis^68^. Structural features were quantified on MG and MT level using featureCounts^97^. Taxonomic annotation of reads and contigs was performed using Kraken2^98^ with a GTDB release207 database (http://ftp.tue.mpg.de/ebio/projects/struo2/GTDB_release207/kraken2) and a 0.5 confidence threshold. Additionally, taxon abundances were estimated using mOTUs 2.5.1^99^. The mOTU abundances were used to generate abundance matrices for each taxonomic rank (phylum, class, order, family, genus and species) by summing up taxon marker read counts at the respective levels.

### Metaproteomics prediction and annotation

For each sample, the predicted proteins were concatenated with a cRAP database of contaminants and the human UniProtKB Reference Proteome prior to the MP search. In addition, reversed sequences of all protein entries were added to the databases for the estimation of false discovery rates. The search was performed using Sipros v1.1^100^ as search engine with the following parameters: trypsin was used as the digestion enzyme and a maximum of two missed cleavages was allowed. The tolerance levels for matching to the database were 1 Da for MS1 and 0.01 Da for MS2. Peptides with large errors for parent ions were later filtered out by setting the Filter Mass Tolerance Parent Ion parameter to 0.05 Da. Carbamidomethylation of cysteine residues was set as a fixed modification and oxidation of methionines was allowed as a variable modification. Peptides with length between 7 and 60 amino acids, with a charge state composed between +2 and +4 and a maximum missed cleavages of 3 were considered for identification. The results from all identifications were filtered by Sipros using at least one unique peptide per protein and peptide false discovery rate (FDR) was dynamically set to achieve a 1% of protein FDR.

Data analysis was performed on all samples with at least 2000 proteins identified. A summary matrix of all selected samples consisting of the KO annotations from the integrated MG and MT analysis and the spectral count from the MP identification was then generated and used for statistical analysis.

## QUANTIFICATION AND STATISTICAL ANALYSIS

### Dimensionality reduction and ordination

Beta diversity for MG and MT was assessed using the Bray-Curtis dissimilarity and subjected to a Non-Metric MultiDimensionnal Scaling (NMDS) for both the taxonomic and functional levels, using the *metaMDS()* function from the vegan package (2.6.2). Principal Component Analysis (PCA) was performed for MP and MM using the *rda()* function in the vegan package (2.6.4). PERMANOVA was used to assess statistical differences between groups using the Bray-Curtis dissimilarity and conducted in the vegan R package with the *adonis2()* function.

### Differential abundance analysis and correlations

Differential abundance was done in two different approaches. The first approach consisted in using SIAMCAT^72^ and ALDEx2^80^ algorithm to find all the taxa and genes differentially expressed between groups without prior assumption. We used two different algorithms to have a sensitive algorithm (SIAMCAT, less prone to have false negative) and a more conservative one (ALDEx2, less prone to have false positive). The second approach consisted in using the MM significant compounds to drive the analysis on the functional level for MG and MT. Therefore, differential abundance tests and multiple correlation tests were conducted with a classical approach. We used Mann-Whitney or Kruskal-Wallis followed by a Dunn test (depending on the number of groups) and Spearman correlation tests. We applied FDR correction using the Benjamini & Hochberg method^101^. We depicted both FDR corrected as q-values and non-FDR corrected p-values to represent most of the differences found in our datasets. All statistical tests were done using the *rstatix* package (0.7.2).

### Variance analysis

Variance analysis was used to assess the importance of each clinical factor on MM. To verify the covariance of factors and to assess which factors explained the most variance in our datasets, we computed the total variance for each clinical factor (removing the NAs for each factor) and the variance explained by each group within a clinical factor. Explained variance was calculated as follows: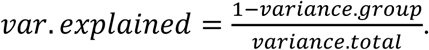

### Microbiome-wide metabolic network analysis

The microbiome-wide metabolic network analysis was conducted by establishing an association between KEGG KOs and corresponding ChEBI IDs. The networks were visualized utilizing the NetworkX package (release 3.3)^73^. In this network, the nodes were represented by KEGG KOs, while the edges were denoted by the corresponding metabolites (either products or reactants)^34^. Such compounds as water, energy transporters and cofactors were removed, to only consider main compounds of a given reaction. The analysis was restricted to genes that were present in a minimum of 50% of the samples. To construct metabolite-specific networks, we used KEGG KOs which have either a reactant or product in KEGG. Glutamate-, thymine-, glycerol-, serine-, alanine-and glucuronate-specific subnetworks were composed of 146, 9, 66, 43, 70, 18 genes, respectively. The network topology metric ‘Betweenness centrality’ was used to underscore the importance of a metabolite in microbiome-wide metabolism^34^.

### Integrated multi-omics analysis using MOFA2

Integrative analysis for the seven omics layers was conducted with the Multi-Omics Factor Analysis (MOFA) 2 R package (version 1.10.0)^102^. Before the analysis, data were preprocessed as follows: (I) funMG, funMT, funMP, taxMG, taxMT, and taxMP data were filtered based on the number of non-zero features, a feature was kept if it was present in at least in 25% of samples in each group (PD, HC, iRBD) or at least in 75% in any of the groups; (II) funMG, funMT, taxMG, and taxMT count data were separately residualized in a linear model to remove variance explained by differences in sequencing depth; (III) funMP and taxMP data were residualized by the sum of protein counts per sample and information on the number of high-quality proteins recovered per sample; (IV) regression residuals were cubic-root transformed to account for heteroscedasticity; (V) MM data were transformed using a centered log-ratio transformation. Each dataset was then additionally filtered to retain the features with the largest variance for the subsequent analysis. For funMG and funMT, we included features with variances equal to or larger than 90% feature variance for a dataset, for the other datasets we included features with variance equal to or larger than the median feature variance for a given dataset. In the results, the feature size for omics layers was as follows: 759 for funMG, 657 for funMT, 410 for funMP, 109 for taxMG, 71 for taxMT, 115 for taxMP, and 34 for MM. MOFA analysis was run on scaled omics data with fifteen initial factors. All factors that explained less than 2% of the variance were excluded from the model. The remaining factors were tested for differential abundance between the groups studied using the linear regression followed by ANOVA type II controlling the participants’ sex, age, and recruitment cohort.

**Extended figure 1.**

NMDS analysis of **A.** metagenomic taxonomic composition, **B.** metagenomic functions and **C.** meta-proteomic taxonomic composition, using a Bray-Curtis dissimilarity matrix. **D.** PCA analysis of metaproteomic functions. **E.** PERMANOVA analysis for the three groups and all omics. Colour represents R² values and size is –log10(p-value). All PERMANOVA analysis were run using 1000 permutations using a Bray-Curtis dissimilarity matrix.

**Extended figure 2.**

**A.** Differential abundance analysis at the genus level using SIAMCAT algorithm. **B.** Differential abundance analysis using ALDEx2 algorithm at the species level. Values are pseudo fold changes for HC/PD and size is based on –log10(p-value). Shape is referring to level of significance, triangular shape for p-value significance before and round shape for p-values < 0.05 after FDR correction.

**Extended figure 3.**

**A.** Percentage of variance explained for each metabolite for “Sex” and “Diagnosis” (left panel) and Percentage of variance explained for each metabolite for “Constipation” and “Diagnosis” (right panel). Metabolites are including untargeted meta-metabolomics, targeted SCFA and targeted bile acids, normalized by sum before merging and variance quantification. **B.** Variance for each metabolite associated to the clinical factors “Diagnosis”, “Sex” and “Constipation”. Metabolites are including untargeted meta-metabolomics, targeted SCFA and targeted bile acids, normalized by sum before merging and variance quantification. **C.** Spearman correlation between metabolites and taxMG species. P-values are FDR corrected. **D.** Absolute log2 fold change between HC and iRBD for funMG and funMT associated to significant compounds. Dots are scaled by the –log10(p-value), colorized and shaped according to p-value significance before (triangle shape) and after FDR correction (round shape).

**Extended figure 4.**

**A to D.** ALDEx2 differential abundance analysis on funMG for HC vs PD (**A.**) and HC vs iRBD (**B.**); funMT for HC vs PD (**C.**) and HC vs iRBD (**D.**). All genes and transcripts are colorized and shaped according to p-value significance before (triangle shape) and after FDR correction (round shape) **E.** Spearman correlation between beta-glutamate relative abundance and funMG-funMT KEGG orthologs related to glutamate species. Only genes with at least one significant correlation are plotted. All p-values are FDR corrected.

**Extended figure 5.**

**A.** Bile acids transcripts found significantly different between the groups. P-values are corrected with FDR. **B.** Flagellar assembly transcripts encoding for extracellular component of the flagella for the genus present in Cluster 2. All tests are Wilcoxon tests.

**Extended figure 6.**

Multiomics variance explained by MOFA factors. **A**. Variance explained by the MOFA factors across different omics layers, total. **B**. Variance explained by the MOFA factors across different omics layers, splitted by factors.

## Data availability

Due to privacy restrictions, the datasets generated by this study are available upon reasonable request by contacting Paul Wilmes.

## Code availability

The IMP pipeline, which was used for analysis of metagenomic and metatranscriptomic data, is available at https://gitlab.lcsb.uni.lu/IMP/imp3. The R and python code used for statistical analyses and visualizations is available at https://gitlab.lcsb.uni.lu/ESB/[TBA].

## Acknowledgments

We thank the staff of the Luxembourg Centre for Systems Biomedicine (LCSB), particularly the sequencing platform and the metabolomics platform, for running the sequencing and metabolomic analyses. The bioinformatics presented in this paper were carried out using the HPC facilities of the University of Luxembourg^103^.

This project has received funding from the European Research Council (ERC) under the European Union’s Horizon 2020 research and innovation programme (grant agreement No. 863664), and was further supported by the Luxembourg National Research Fund (FNR) CORE/16/BM/11333923 (MiBiPa), CORE/15/BM/10404093 (microCancer/MUST), PRIDE/11823097 (MICROH DTU), and CORE/19/BM/13684739 (metaPUF), the Michael J. Fox Foundation under grant IDs 14701 (MiBiPa-PLUS) and MJFF-019228 (PARKdiet), the Parkinson’s Foundation (MiBiPa Saliva), the Institute for Advanced Studies of the University of Luxembourg through an AUDACITY grant (ref. no. MCI-BIOME_2019), as well as an “Espoîr en tête” grant from the Rotary Club Luxembourg to P.W. This work was also supported by a Fulbright Research Scholarship from the Commission for Educational Exchange between the United States, Belgium and Luxembourg to P.W. Additional funding was provided by the FNR under INTERMOBILITY/23/17856242. C.d.R. received funding from the European Union’s Horizon 2020 Widening Fellowships (N° 101026381).

## Contributions

Conceptualization: J.-P.T., A.H.-B., W.O., B.M., P.W. Patient recruitment, clinical coordination, and sampling: S.S., A.J., C.T., W.O., B.M. Multi-omic data generation: C.D.R., J.-P.T., C.J., L.A.L, A.D, N.J, M.v.B. Bioinformatics, statistics and data visualization: R.V, J.O.S., P.N., V.T.E.A, V.P, O.H, B.K, P.M. Initial manuscript draft: R.V, J.O.S, P.N, V.T.E.A, V.P. Review and editing: C.C.L., S.B.B., P.M. and P.W. Funding acquisition: C.C.L, W.O, B.M, P.W. All authors read and approved of the submitted version.

## Rights Retention Statement

This research was funded in whole, or in part, by the Luxembourg National Research Fund (FNR), grant references CORE/16/BM/11333923 (MiBiPa), CORE/15/BM/10404093 (microCancer/MUST), PRIDE/11823097 (MICROH DTU), and CORE/19/BM/13684739 (metaPUF). For the purpose of open access, and in fulfilment of the obligations arising from the grant agreement, the author has applied a Creative Commons Attribution 4.0 International (CC BY 4.0) license to any Author Accepted Manuscript version arising from this submission.

## Extended information

### Multi-omics data overview

Using our previously developed methodological framework^107,108^, we performed a systematic multi-omic analysis of DNA, RNA, protein, and metabolite fractions isolated from flash-frozen fecal samples. We used MG, MT, MP and MM data to find biomarkers associated with the PD phenotype (Fig. 1). We generated a mean of 7.5 (std 1.7) Gbps and 7.5 (std 1.4) Gbps of sequencing data for MG and MT, respectively. After trimming and filtering, we retained a mean of 6.8 (std 1.7) Gbps and 3.2 (std 1.3) Gbps for MG and MT, respectively. The mean assembly size was 0.4 (std 0.1) Gbps, with on average 5.9×10^5^ (std 1.7×10^5^) genes predicted. Finally, protein databases contained a mean of 7.2×10^5^ (std 1.8×10^5^) proteins, an average of 4.1×10^4^ (std 0.6×10^4^) MS spectra per sample were acquired, and a mean of 3.4×10^3^ (std 1.7×10^3^) proteins were identified.

### MOFA model description

MOFA is an unsupervised machine learning approach for the integration of multi-omics data sets^102^. It allows for the identification of highly informative features across multiple omics. It has previously been used in the study of the gut microbiome in several diseases, giving critical insights into the link between the gut microbiome, health, and disease^104–106^. The biggest proportion of variance was explained by funMG and funMT, followed by the taxMT and funMP datasets (Extended fig. Aa). F1-2 incorporated most of the variance related to the funMT and taxMT, whereas funMG and taxMG variance was predominantly covered by F3- F5 (Extended fig. 8B). The funMP variance was explained mostly by F6, and MM variance was explained by F1 and F6. MOFA factors were tested in a linear model followed by ANOVA with disease status, as well as confounders including patients’ sex, age, and recruitment cohort.

Among the MOFA factors, F1 showed an association with the disease status, whereas F4 and F9 were associated with patients’ sex and recruitment cohort, respectively (Fig. 4A).

