## Supplementary figures and images for "Integrated multi-omics highlights alterations of gut microbiome functions in prodromal and idiopathic Parkinson’s disease"

### Extended figure 1

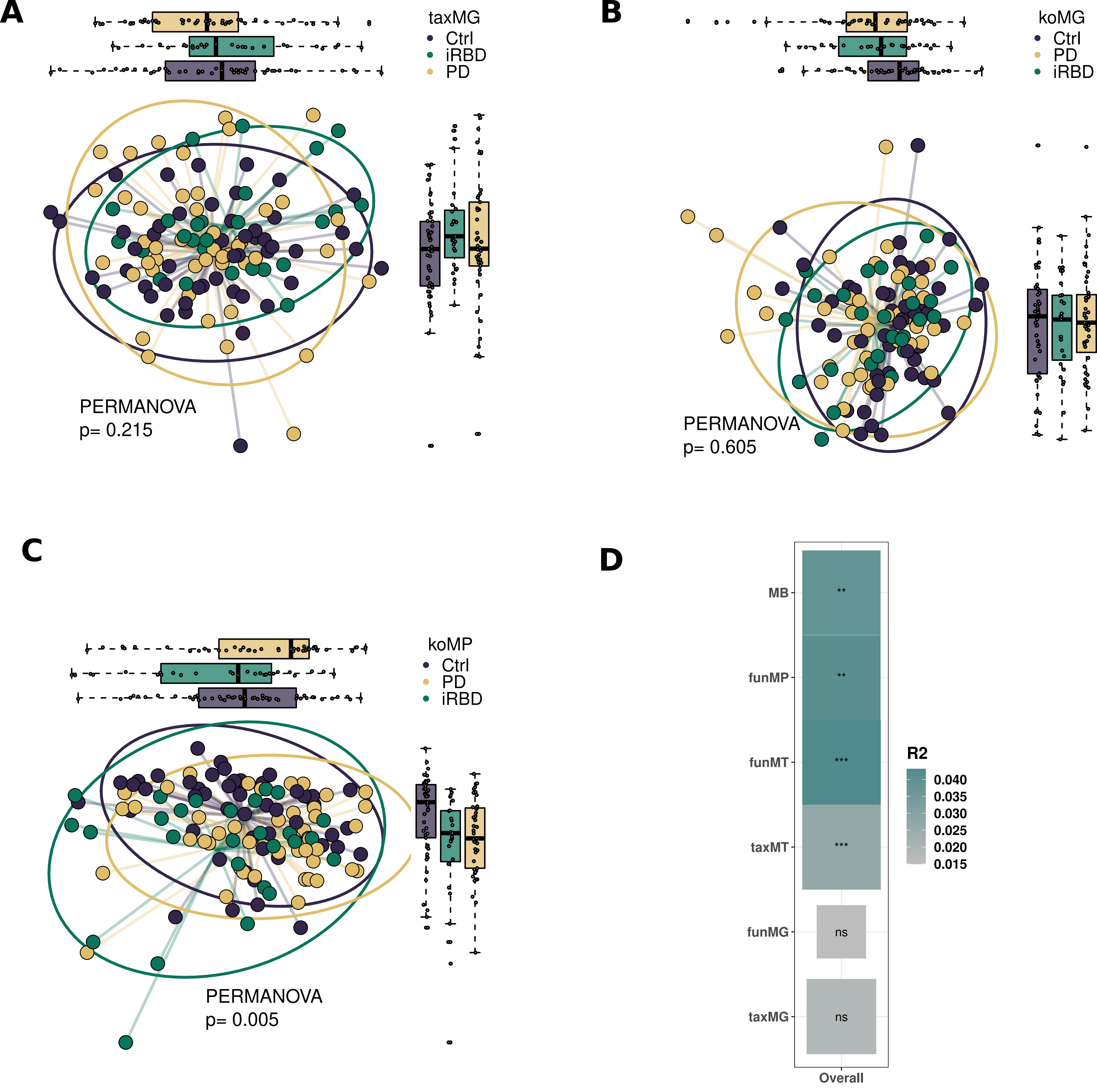

### Extended figure 2

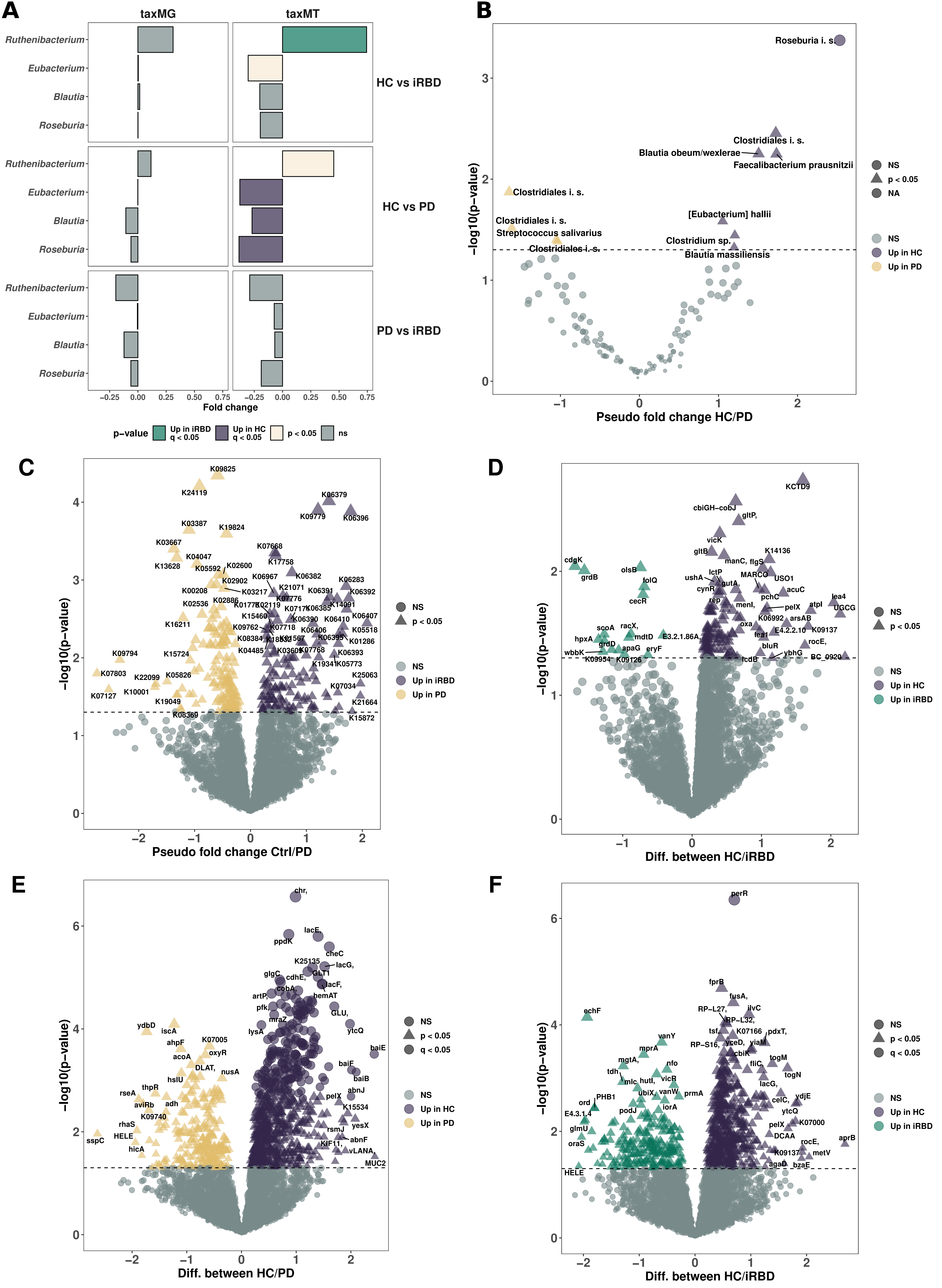

### Extended figure 3

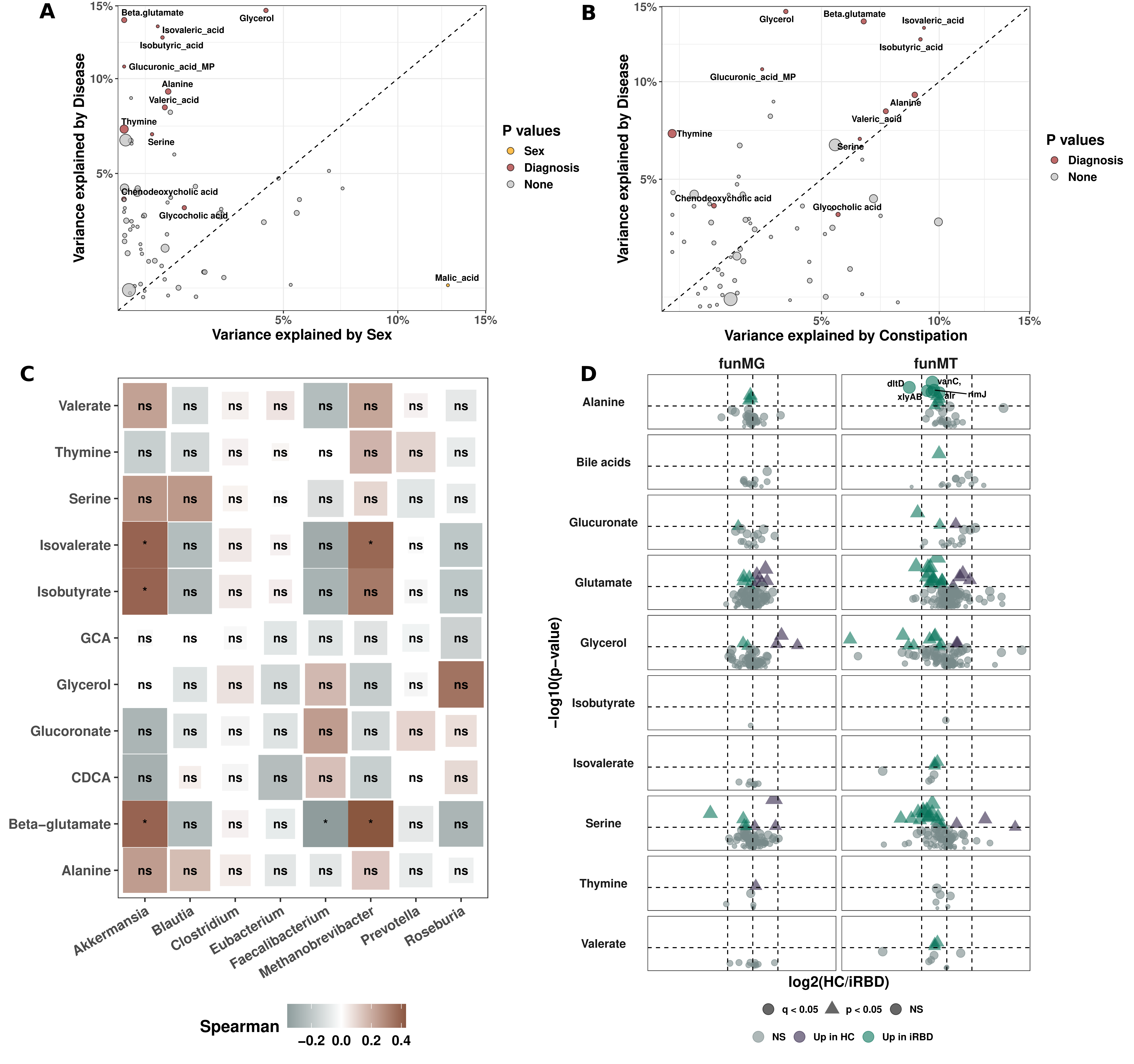

### Extended figure 4

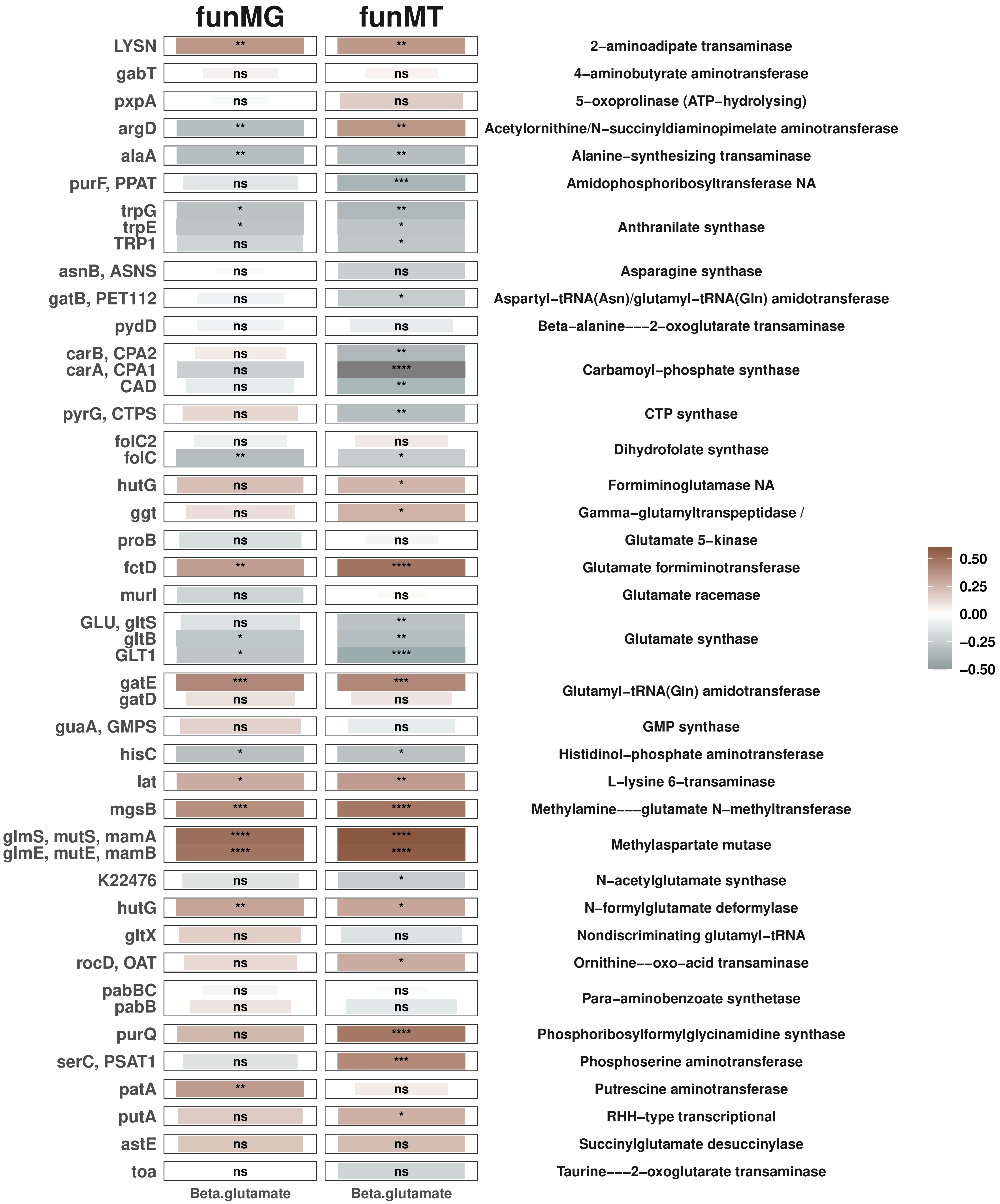

### Extended figure 5

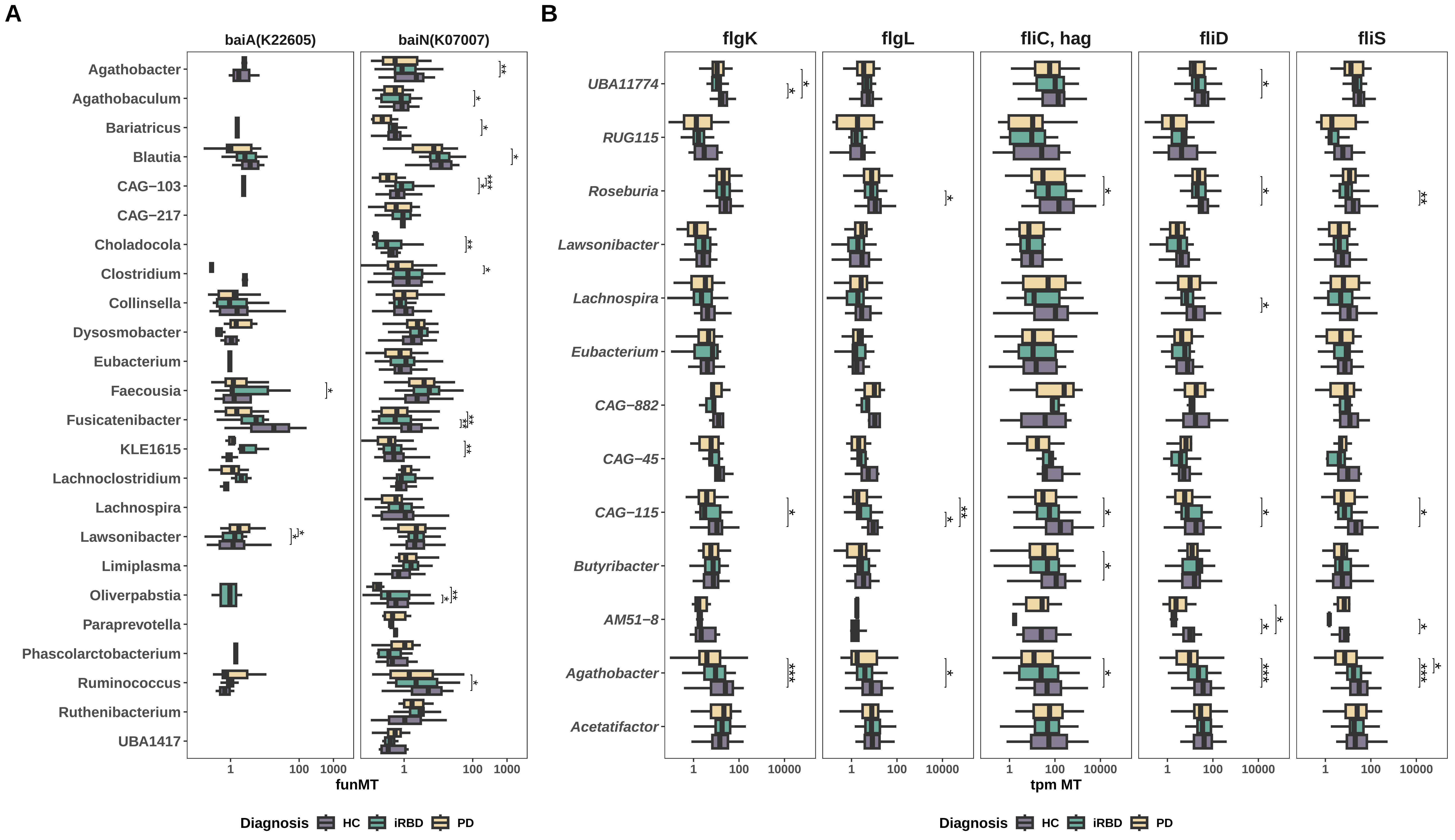

### Extended figure 6

Multiomics variance explained by MoFa factors

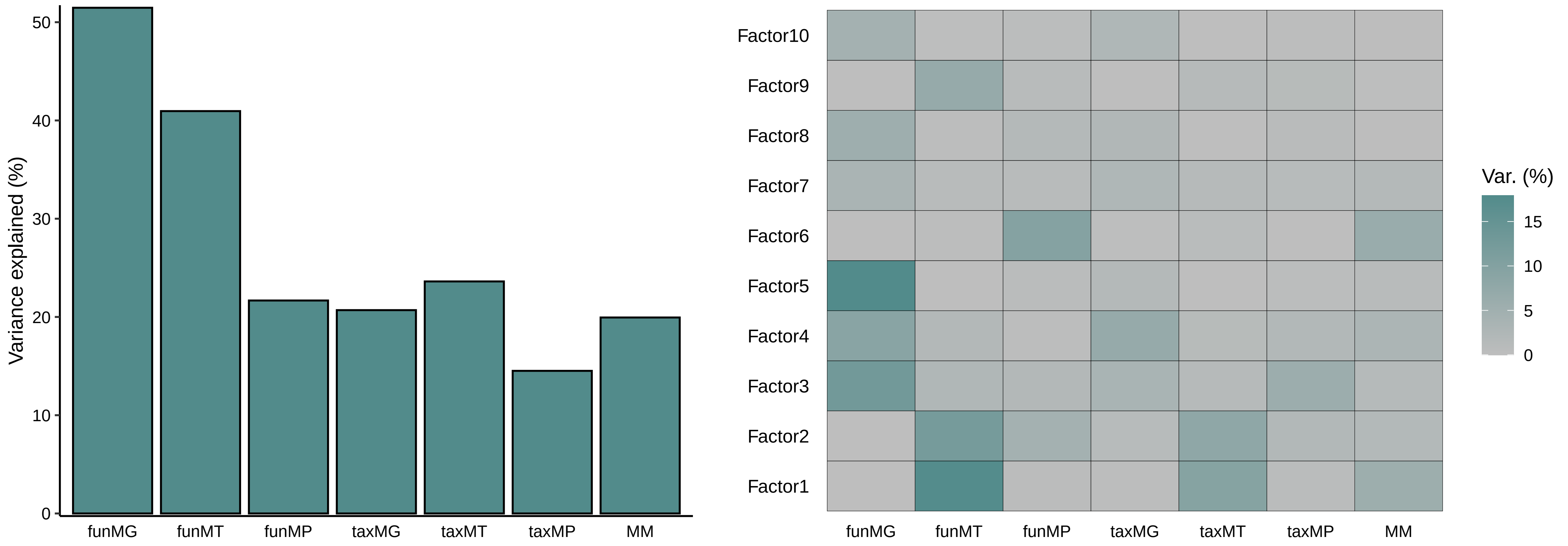
